# SETD3 ACTIN HISTIDINE METHYLTRANSFERASE REGULATES INTESTINAL EPITHELIAL BARRIER INTEGRITY AND MUCOSAL INFLAMMATION

**DOI:** 10.64898/2026.10.02.756302

**Authors:** Prasath Manogaran, Nayden G. Naydenov, Susana Lechuga, Naseer Sangwan, Andrew Deng, Ajay Zalavadia, Andrei I. Ivanov

## Abstract

Intestinal epithelial barrier is a key feature of the healthy gut, and disruption of this barrier is a driver of mucosal inflammation. Integrity of the intestinal epithelial barrier is controlled by tight junctions (TJs) and adherens junctions (AJs) associated with the underlying actin cytoskeleton. Posttranslational modifications of actins affect the cytoskeletal assembly and dynamics, however whether such modifications regulate epithelial barriers remains unknown. SETD3 methyltransferase methylates actins at the His73 residue, thereby enhancing filamentous (F)-actin polymerization and turnover. This study examines the role of SETD3 in regulating structure and function of normal intestinal epithelial barrier, as well as its involvement in barrier disruption during mucosal inflammation. siRNA mediated knockdown of SETD3 in DLD-1 and HCA-7 human intestinal epithelial cells (IEC) resulted in several functional and morphological cell abnormalities. These include increased permeability of the IEC barrier, diminished assembly of the perijunctional F-actin belt, increased cell size, and delayed TJ reassembly during calcium switch. Consistently, SETD3 knockout (KO) mice displayed increased intestinal permeability and decreased labeling intensity of apical β-actin as compared to wildtype controls. Furthermore, SETD3 KO mice showed increased cell stemness and proliferation in colonic epithelium without defects in AJ/TJ assembly and secretory cell differentiation. In addition, SETD3 KO mice developed more severe disease symptoms during dextran sodium sulfate (DSS)-induced colitis, which were accompanied by accelerated mucosal erosion and increased macrophage, T cell and neutrophil infiltration into colonic tissue. Overall, our data suggests that SETD3 acts as a novel positive regulator of the intestinal epithelial barrier and plays a protective role during intestinal mucosal damage and inflammation.

## Introduction

Establishment of the intestinal epithelial barrier plays multiple homeostatic and protective roles, such as regulated transepithelial exchange of different substances and the establishment of symbiotic relationships between the host and the gut microbiota ^1–3^. Disruption of the gut barrier is a common manifestation of various intestinal and extraintestinal diseases that perpetuate and sustain inflammatory processes throughout the body ^1, 4–10^. Gut barrier integrity is mediated by the assembly of several junctional complexes located at epithelial cell-cell contacts. Tight junctions (TJs) and adherens junctions (AJs) are the major determinants of intestinal barrier integrity and permeability ^11–16^. The molecular composition of TJs and AJs consists of adhesive transmembrane proteins, whose spatial position at the cell surface is controlled by elaborate scaffolding platforms assembled on the cytosolic face of the plasma membrane ^11–16^. Tetraspan proteins such as occludin and claudins, along with Ig-like proteins such as Junction adhesion molecule A are the most well-known transmembrane TJ components that interact with multiple types of cytoplasmic scaffolds, most notably with members of the ‘*zonula occludens*’ (ZO) protein family ^11, 12, 15^. Similarly, E-cadherin is a calcium-dependent, adhesive transmembrane AJ protein in the intestinal epithelium that interact with a scaffolding cytoplasmic complex that consists of α-catenin, β-catenin and p120-catenin ^13, 14, 16^.

Assembly and remodeling of intestinal epithelial apical junctions is tightly regulated by several mechanisms that target the expression, stability and localization of junctional proteins ^5, 11, 12, 17^. One of the most important regulatory mechanisms involves direct coupling of AJs and TJs to the underlying cortical actin cytoskeleton ^12, 18–23^. Differentiated, barrier-forming intestinal epithelial cells are characterized by a prominent apical circumferential actin cytoskeletal belt attached to AJs and the basal part of TJs via different actin-binding proteins presented in the junctional cytoplasmic plaques ^12, 18–23^. This circumferential actin belt is essential for clustering and stabilization of junctional complexes. Furthermore, this belt is enriched with non-muscle myosin II (NM II) motors that generate mechanical forces critical for the stability and remodeling of epithelial cell-cell contacts ^3, 12, 18, 19, 23, 24^.

Junctional associated actin filaments are composed of two different actin paralogs, β-cytoplasmic and γ-cytoplasmic actins, that have unique and redundant functions in regulating epithelial barrier integrity *in vitro* and *in vivo* ^25, 26^. Assembly, turnover and bundling of these filaments is controlled by a variety of actin-nucleating, capping, cross-linking and depolymerizing proteins, as highlighted by the studies examining the molecular composition and interactomes of AJs and TJs ^27–30^. Furthermore, a large body of evidence supports the causal roles of several actin-binding proteins in controlling epithelial junctional assembly and remodeling ^31–40^.

Much less is known about whether the structural properties of actin molecules could regulate cytoskeleton-driven remodeling of apical junctions and the establishment of epithelial barriers. Importantly, actin filament assembly and dynamics depend on various post-translational modifications (PTMs) of the actin molecules, most notably acetylation, methylation, arginylation, and oxidation^41, 42^. These PTMs regulate actin filament polymerization and interactions with different accessory proteins and have been shown to regulate different cellular processes including cell division, migration, and differentiation of specialized neuronal and muscle cells ^41, 42^. Yet, it is unclear whether actin PTMs are important towards the assembly of apical junctions and the establishment of intestinal epithelial barriers.

SET domain 3 (SETD3)-mediated actin methylation is an emerging regulator of the actin cytoskeleton. Methylation of actin molecules at the His73 residue is one of the best-known actin modifications that has been discovered more than 50 years ago^43, 44^. Biochemical studies in cell-free systems demonstrated that His73 methylation diminishes the rate of actin polymerization, affects ATP hydrolysis and decreased thermal stability of monomeric actin ^45–48^. However, the biological relevance of this actin modification only began to emerge after the discovery of SETD3-dependent His73 actin methylation in cells and live organisms ^45, 48^. SETD3 was originally reported to methylate Histone 3 at Lysines 4 and 36^49^. However, this conclusion was challenged by subsequent studies which identified SETD3 as a *bona fide* histidine methyltransferase that does not modify histones but catalyzes a His73 methylation of mammalian actins^45, 48^. Importantly, actin appears to be a major, if not the only, substrate of SETD3 in mammalian cells^45, 48^, as depletion of SETD3 resulted in almost complete loss of actin His73 methylation^45, 48^. Biochemical and imaging studies reveal that the loss of SETD3 resulted in abnormal organization and decreased polymerization of the actin cytoskeleton in cancer and neuronal cells^45, 50–52^. Furthermore, functional studies of SETD3 reveal that this methyltransferase mediates cell proliferation and migration in liver and breast cancer cells^51, 53–55^. However, nothing is known about the role of SETD3 in regulating the integrity and remodeling of epithelial barriers under homeostatic or inflammatory conditions. The present study is designed to close this knowledge gap and investigate SETD3-dependent regulation of the intestinal epithelial barrier integrity both *in vitro* and *in vivo*. We report that SETD3 promotes the assembly of apical junctions and the establishment of the paracellular barrier in model IEC monolayers by controlling integrity of junction-associated actomyosin cytoskeleton. Furthermore, our findings suggest that SETD3 enhances permeability of normal gut barrier and plays a protective role during mucosal inflammation *in vivo*.

## Material Methods

### Antibodies and other reagents

The following antibodies were used for detecting cytoskeletal, junctional, and leukocyte proteins: SETD3 (Cell Signaling, Danvers, MA, Cat# 99124 & ABclonal Science, Woburn, MA Cat# A8071), ZO-1 (Thermo Fisher Scientific, Waltham, MA, Cat# 33-9100 (immunoblotting) and BiCell Scientific, Maryland Heights, MO, Cat# 00235 (immunofluorescence)), E-cadherin (R & D Systems Minneapolis, MN, Cat# AF748), occludin (BiCell Scientific, Cat# 00241), β-catenin (BD Biosciences, San Jose, CA, Cat# 610154,), α-catenin (BD Biosciences, Cat# 610194), claudin-3 (Millipore-Sigma, Burlington, MA, Cat# SAB4500434), claudin-4 (Thermo Fisher Scientific, Cat# 32-9400), claudin-2 (Thermo Fisher Scientific, Cat# 51-6100), claudin-7 (Thermo Fisher Scientific, 34-9100), NM IIC (Cell Signaling, Cat# 8189S), NM IIA (BioLegend, San Diego, CA, Cat# 909801), total MLC (Cell Signaling, Cat# 3672S), GAPDH (Cell Signaling, Cat# 2118S), total actin (Millipore-Sigma, Cat# MABT219), ppMLC (Cell Signaling, Cat# 3674S), MUC-2 (Abcam, Waltham, MA, Cat# ab272692), lysozyme (Abcam, Cat# EPR2994), DCLK1 (Cell Signaling Technology, Cat # 62257S), Ki-67 (Abcam, Cat# ab15580), OLMF4 (Cell Signaling Technology, Cat # 39141), CD3 (Cell Signaling Technology, Cat # 78588), F4/80 (Cell Signaling Technology, Cat # 70076), Ly6G (Bio legend, Cat# 127601). Mouse monoclonal antibodies selectively recognizing β-cytoplasmic actin and γ-cytoplasmic actin we provided by Dr. Christine Chaponnier (University of Geneva, Switzerland) ^25^. A rabbit polyclonal antibody that recognizes His73-methylated actin was provided by Dr. Or Gozani (Stanford University School of Medicine) ^48^. Alexa Fluor 488 and Alexa Fluor 555–conjugated donkey anti-rabbit, donkey anti-goat and donkey anti-mouse secondary antibodies as well as Alexa Fluor-488-conjugated Phalloidin were obtained from Thermo Fisher Scientific. Horseradish peroxidase–conjugated goat anti-rabbit and anti-mouse secondary antibodies were acquired from Bio-Rad Laboratories (Hercules, CA). SPY650-FastAct-X probe was obtained from Cytoskeleton Inc (Denver, CO, Cat# CY-SC502). All other chemicals were obtained from Thermo Fisher Scientific and Millipore-Sigma.

### Cell culture

DLD-1 cells were purchased from the American Type Collection (ATCC, Manassas, VA, Cat# CCL-221) and HCA-7 colony 29 cells were purchased from Millipore-Sigma, (Cat# 02091238). Both cell lines were found to be mycoplasma free as per Universal Mycoplasma Detection Kit (ATCC, Cat# 30-1012K) and were used up to 10 passages. HCA-7 cells were cultured in high glucose DMEM with low bicarbonate (ATCC, Cat# 30–2002) supplemented with 10% fetal bovine serum (FBS), HEPES, non-essential amino acids, and penicillin-streptomycin antibiotic cocktail. DLD-1 cells were cultured in DMEM/F12 (Corning, New York, NY, Cat#10-092CV) and supplemented with 5% FBS and penicillin-streptomycin antibiotic cocktail. To measure Transepithelial Electrical Resistance (TEER), cells were plated on collagen coated transwell membrane filters. For immunolabelling experiments, cells were seeded on collagen coated glass coverslips. For biochemical studies, the cells were seeded and cultured in six well plates.

### Animals

SETD3 null mice on the C57BL/6N background were described previously ^48^. This strain was initially developed in the Toronto Centre for Phenogenomics and breeding pair were provided by Drs. Jan Carrette and Or Gozani from Stanford University School of Medicine. Heterozygous SETD3 mice were used as breeders and SETD3 null and wild type littermates were used in the experiments. Approximately equal numbers of male and female mice were included in each experimental group, but we did not specifically investigate sex-specific differences in their responses. The animal colonies were maintained under pathogen-free conditions in the vivarium of the Cleveland Clinic Research. Standard feed and tap water were available, *ad libitum*. The mouse room was on a 12 h light/dark cycle (lights on at 7:00 A.M.). All procedures were conducted under an animal research protocol approved by the Cleveland Clinic Research (IACUC protocol # 00001872) in accordance with the National Institutes of Health Animal Care and Use Guidelines.

### Small-interference RNA (siRNA)-mediated knockdown of SETD3

siRNA mediated transient knockdown of SETD3 in DLD-1, and HCA-7 cells was performed using ON-TARGET plus siRNA Smart pool purchased from Horizon Discovery (Water beach, United Kingdom, Cat# L2-014808-01-0005). Scrambled siRNA sequence, which lacks complementarity to any human gene, was used as control siRNA (Horizon Discovery, Cat# D-001810-10-05). Both DLD-1 and HCA-7 cells were transfected using DharmaFECT 1 (Horizon Discovery, Cat# T-2001–02) in Opti-MEM I medium (Thermo Fisher Scientific, Cat# 31985070), according to the manufacturer’s protocol, with a final siRNA concentration of either 50 or 100 nM. All cells were used for experiments on days 3 and 4 post transfection.

### Measurement of epithelial barrier permeability

TEER and FITC-dextran flux measurements in DLD-1 and HCA-7 cells were measured as described previously^33, 34, 36^ with slight modifications. Specifically, 6 × 10^4^ DLD-1 or HCA-7 cells were seeded in collagen coated 24 well transwell inserts (Corning, Cat# 353095) with polyethylene terephthalate membrane filters (0.4 µm pore size). TEER was determined using a EVOM2 VoltOhm Meter (World Precision Instruments, Sarasota, FL). The resistance values were corrected by subtracting a cell-free blank transwell insert from each experimental point and multiplied by the area to obtain final TEER values. For FITC-dextran flux measurement, DLD-1 and HCA-7 cells were cultured on 24 well collagen-coated transwell inserts. 4 days post-transfection, cells were apically exposed to 1 mg/mL FITC-labeled dextran (4,000 Da) in HEPES-buffered Hanks’ Balanced Salt Solution with calcium and magnesium (HBSS^+^). After 4h of incubation, HBSS were collected from the lower chamber, and FITC fluorescence intensity was measured using a BioTek Synergy H1 microplate reader (Agilent Technologies), at excitation and emission wavelengths 495 nm and 525 nm, respectively. The amount of translocated FITC-dextran across the epithelial cell monolayer was calculated based on a calibration curve generated using Prism 10 (GraphPad, La Jolla, CA).

### Immunoblotting analysis

On day 4 post transfection, DLD-1 & HCA-7 cells were scraped in RIPA buffer (20 mM Tris, 50 mM NaCl, 2 mM EDTA, 2 mM EGTA, 1% sodium deoxycholate, 1% Triton X-100 (TX-100), and 0.1% SDS, pH 7.4) supplemented with protease inhibitor cocktail, phosphatase inhibitor cocktails 2 and 3, and Pefabloc^®^ (Millipore-Sigma, Cat# P8340, P5726, P0044, and 76307 respectively). Cells were then homogenized via Dounce homogenizers, and lysates were centrifuged at 14,000 x g for 20 min. Supernatants were then diluted with 2x Laemmli sample loading buffer, supplemented with 2 mM DTT, and boiled at 95 C for 5 minutes.

Samples were then processed using SDS-polyacrylamide gel electrophoresis, for which either 10 μg or 20 μg total protein were loaded per lane. The separated proteins were then transferred to nitrocellulose membranes, membranes were blocked with 5% non-fat milk for 1 hour, incubated with 1:1,000 diluted primary antibody overnight, and then incubated with 1:5,000 diluted HRP-conjugated secondary antibodies for 1 hour. Membranes were then submerged in enhanced chemiluminescence solution (Millipore Sigma, GERPN2209), and proteins were visualized through X-ray film exposure. Band intensities were quantified using the gel densitometry toolset from ImageJ 1.51K software (National Institute of Health, Bethesda, MD).

### Calcium switch assay

The calcium switch assay was carried out as we described previously ^25, 33, 34, 36, 56^. Briefly, on day-3 post-transfection, DLD-1 & HCA-7 cells monolayers were subjected to overnight extracellular calcium depletion by replacing the media with Eagle’s minimum essential medium for suspension culture (Gibco, Cat# 11380037) supplemented with 5 µM CaCl_2_, 10 mM HEPES and 10% dialyzed FBS, pH 7.4. Cells were then incubated with their original cell culture medium, which contains high calcium, at the indicated times to facilitate junctional reassembly.

### Immunofluorescence labeling, confocal microscopy, and image analysis

To determine the organization of epithelial junctions, as well as SETD3 and NMIIA localization DLD-1 and HCA-7 cells were fixed and permeabilized with 100% ice-cold ethanol or methanol for 20 min at −20°C. To detect di-phosphorylated myosin light chain (ppMLC) and F-actin, cells were fixed with 4% paraformaldehyde (PFA) for 20 min at room temperature, then permeabilized with 0.5% Triton X-100 in HBSS^+^ for 5 min at room temperature. Following fixation and permeabilization, cells were blocked with 1% bovine serum albumin (BSA), pH 7.4 for 1 hour and incubated with primary antibodies diluted 1:200 in blocking buffer for 1 hour at room temperature. Cells were then washed thrice with 1% BSA and were incubated with Alexa-Fluor-488–conjugated donkey anti-rabbit and Alexa-Fluor-555–conjugated donkey anti-mouse secondary antibodies diluted 1:1000 with blocking buffer for 1 hour at room temperature. Lastly, cells were washed three more times with blocking buffer and mounted on slides with ProLong™ Gold Antifade mounting medium supplemented with DAPI (Thermo Fisher Scientific, Cat # P36941). For F-Actin labeling, fixed/permeabilized cells were incubated for 1 hour with Alexa-Fluor-488 conjugated phalloidin diluted 1:1000 in blocking buffer, washed 3 times in blocking buffer, and then mounted with mounting media. Immunofluorescence-labeled cell monolayers were imaged using a Leica SP8 confocal microscope (Wentzler, Germany). The Alexa Fluor 488 and 555 signals were acquired sequentially in frame-interlace mode, to eliminate cross-talk between channels.

In the calcium-switch experiments, fluorescence signal intensities for adherens junction tight junction, and cytoskeletal markers were quantified using QuPath (QuPath v0.7.0, Open Software for Bioimage Analysis). Junction and cytoskeletal reassembly quantification were performed as follows: in each image, the junctional lengths were measured in the entire image and normalized by number of cells. Measurements were obtained from three images per experiment across three independent experiments. To measure junctional accumulation of F-actin and NM II in Fig. 4 images, the ImageJ MEAN INTENSITY function was used. This function calculates the arithmetic average of all pixel intensities within the selected junctional region. The data shown are from three images per experimental group obtained in three independent experiments.

Cell size and cell height in control and SETD3-depleted DLD-1 and HCA-7 cell monolayers were measured using the CPSAM extension in QuPath. Data was collected from three independent experiments, with three images analyzed per group in each experiment.

### Live cell imaging and data analysis

Junctional actin dynamics was assessed using the SPY650-FastAct-X probe, according to the manufacturer’s protocol. Briefly, DLD-1 & HCA-7 cells were transfected on collagen-coated 35mm coverslip bottom dish. On day 4 post transfection, the cells were incubated with working concentration (1x in DMSO) SPY650-FastAct-X diluted 1:500 in respective cell culture medium and incubated for 2 hours before imaging. Afterward, the probe was removed by washing the cells with fresh cell culture medium. Live cell imaging was performed with Leica TCS SP8 AOBS confocal laser scanning system on a Leica DMi8 inverted microscope using a Leica 63x CS2 (1.4NA) oil immersion objective and a stage top incubator (OKOLAB United States Inc. Sewickley, United States) with temperature at 37°C and 5% CO2 concentration. 0.70% of 633nm of available laser intensity and a moderate pixel dwell time of 0.4 µs was used for image acquisition with 1 minute interval to limit the phototoxicity. QuPath v0.7.0 was used to measure F-actin intensity over time in both control siRNA and SETD3 siRNA knockdown cells.

### Measuring intestinal permeability in vivo

Intestinal permeability of unchallenged SETD3 null mice and their wild type littermates was examined as we previously described^26, 57, 58^. Briefly, animals were subjected to overnight food deprivation and then were given a solution of FITC-labeled dextran (4 kDa; 80 mg/100 g body weight) via oral gavage. After 3 hours, animals were euthanized, and blood was collected through cardiac puncture. Blood serum was separated by centrifugation, and FITC-dextran fluorescence intensity in the serum samples was measured using a Bio Tek Synergy H1 plate reader (Agilent Technology) with excitation and emission wavelengths at 495/525nm. The fluorescence intensity of dextran-free serum was subtracted from each measurement. The concentration of FITC-labeled dextran in blood serum was calculated using standard curves generated via serial dilutions of a FITC-dextran stock solution in phosphate buffered saline. In order to compare results from different experiments, we calculated a relative FITC-dextran permeability by normalizing the obtained FITS dextran concentration values in each animal sample to a value of one randomly selected wild-type control sample for each experiment.

### Characterization of experimental colitis in SETD3 null mice

Dextran sulfate sodium (DSS) colitis was induced in 8–10-week-old SETD3 null mice and their wild-type control littermates by administering a 2.5% (w/v) solution of DSS (molecular weight range 35,000-50,000 kDa; Thermo Fisher, Cat# J14489.22) in drinking water. Unchallenged animals received tap water. Animals were weighed daily and monitored for signs of intestinal inflammation. The disease activity index was calculated as previously described by averaging numerical scores of body weight loss, stool consistency, and intestinal bleeding ^57, 58^. On day 7 of DSS administration, animals were euthanized, their intestinal tissue harvested and separated into several segments. The samples were either fixed in a 4% formaldehyde solution or embedded in an optimal cutting temperature medium and snap-frozen for subsequent histological and biochemical examination. Formalin-fixed samples were paraffin-embedded, sectioned, and stained with hematoxylin and eosin (H&E). The tissue injury index was calculated based on microscopic examination of H&E sections, as previously described ^59^. The H&E-stained slides were scored in a blinded fashion by a gastrointestinal pathologist. The index represents the sum of individual scores reflecting severity and extent of inflammation, extent of crypt damage and percentage of the involved area.

### Fluorescence labeling of mouse intestinal tissues

Harvested colonic and ileal tissue samples of SETD3 null mice and their wild type controls were paraffin-embedded and sectioned at 5μm thickness. Tissue sections were deparaffinized, subjected to antigen retrieval under the following conditions: sodium citrate pH 6.0 for DCLK1, OLMF4, Lysozyme, E-cadherin, β-catenin and CD3 antibodies; universal antigen retravel buffer, pH 8.0 for Ki-67, MUC-2, β-Actin, γ-Actin, E-cadherin, and β-catenin antibodies; Tris-EDTA pH 9.0 for F4/80, Ly6G, occludin, and ZO1 antibodies, following by blocking for 60 min in a protein free blocking solution (Agilent Technology, Cat# X090930-2). Sections were then stained with primary antibodies diluted in antibody diluting solution (DAKO, Cat# S080983-2) overnight followed by 1 h incubation with secondary antibodies diluted with the antibody diluent solution. Immunolabeled sections were then washed three times with PBS, incubated with Alexa dye conjugated secondary antibodies for 60 minutes, washed three times with PBS, and mounted on slides with ProLong™ Gold Antifade mounting medium with DAPI. Immunofluorescence labeling of Claudin 3 was performed on frozen sections of mouse colonic mucosa. Sections were fixed with ice cold absolute methanol at -20°C for 5 minutes and immunolabeled as described above.

Immunofluorescence labeled tissues were imaged using Leica 63x CS2 (1.4NA) OIL immersion objective equipped on a Leica TCS SP8 AOBS confocal laser scanning system attached to a Leica DMi8 inverted epifluorescence microscope (Wentzler, Germany).

Quantification of proliferative cells (Ki67), intensity of lysozyme, apical junctional β-actin and γ-actin intensity in control and SETD3 KO mice was performed by using Image J. Small rectangular boxes were manually drawn over the apical epithelial surface, and the fluorescence intensity of either β-actin or γ-actin within the selected region was measured using ImageJ and for the quantification of number of Ki67 cells and intensity of lysozyme in the crypt area, small circle were made on the crypt cells and measure the intensity of lysozyme or number of ki67 positive cells per crypt. Three different images per tissue section were quantified for each animal and the obtained values were averaged to produce a single data point.

Quantification of stem cells marker (OLMF4), secretary cells (MUC2 & DLCK) and infiltrated inflammatory cells (F4/80, CD3, Ly6G) was performed using QuPath v0.7.0. Whole-microscopic images acquired at ×63 magnification were imported into QuPath, and positive cells were detected based on the summed optical density in the green channel. For each marker, positive cells were quantified in three images, and the values were averaged to produce a single data point for each animal.

### Bulk RNA sequencing of mouse intestinal epithelial cell samples

IEC isolation was performed as we recently described ^58^. In brief, colonic and ileal epithelial cells were isolated from unchallenged SETD3 KO mice and their wild type littermates by the sequential dithiothreitol/EDTA extraction protocol. Total RNA was extracted using the Qiagen RNeasy Plus Mini kit (Qiagen, Cat#74134), using a modified version of the manufacturer’s protocol. Sample lysates were added to QIAShredder columns and centrifuged, and the homogenized lysates were added to gDNA eliminator columns. The columns were centrifuged, and the lysates were combined with ethanol; the samples were then added to RNeasy Plus Mini spin columns and centrifuged. The membranes were washed with RPE buffer and RNA was eluted using 25 µl of nuclease-free water. The extracted RNA concentrations and the 260/280 absorption ratio were measured using the Qubit assays and Nanodrop. The Total RNA libraries were prepared in the LRI Genomic Core following the manufacturer’s protocol (Illumina, Document #1000000124514). Specifically, ribosomal RNA was depleted, and the RNA underwent fragmentation and denaturation. Subsequently, cDNA was synthesized, followed by end repair, adenylation, and adaptor ligation, culminating in PCR amplification. The libraries were evaluated for quality, quantified, and sequenced at 20 million reads per sample on the Illumina sequencer Novaseq 6000.

### Bioinformatic analysis of bulk RNAseq data

We assessed the read quality with FastQC (v0.12.1[SN1.1^60^, then quality-trimmed the reads and removed adapter sequences using Trimmomatic (v0.39) ^61^ with a minimum quality score of 20 and a sliding window of 4 bases, and rechecked quality with FastQC to confirm the reads met acceptable thresholds. Host contamination was removed by aligning the trimmed reads to the mouse reference genome GRCm39 (Mouse Genome Reference Consortium, Version 39) with BBMap (v38.99) and retaining only non-host reads via the filterbyname option. The cleaned reads were mapped to the mouse genome using STAR (v2.7.10.A) ^62^ with two-pass mapping to improve splice-site detection and reduce multi-mapping reads, producing BAM files from which gene-level counts were generated with feature Counts (Subread package v2.0.1) ^63^ and the corresponding GTF annotation file. Differential expression analysis was performed in R with DESeq2 (v1.38) ^64^, normalizing counts by the median-of-ratios method and applying a negative binomial generalized linear model to identify differentially expressed genes (adjusted p-value < 0.05 and fold change ≥ 2). Pathway enrichment of the DEGs was carried out with enrichR ^65^ against the GO, KEGG, and Reactome databases (*p* value < 0.05), and results were visualized with ggplot2 (v3.4.2) ^66^ as heatmaps of normalized counts for the top DEGs, volcano plots of significance and magnitude of change, and plots of the enriched pathways. The sequence data were uploaded to the NCBI GEO database. GEO ID: GSE344552.

### Statistical analysis

The data is expressed as mean ± standard error (SEM). Statistical analysis was performed using a two-tailed paired test for cell culture experiments (or) unpaired Student’s t-test to compare the results between two data sets for animal experiments. Multi group comparison was performed using one-way ANOVA. *p* values < 0.05 were considered statistically significant and all statistical analysis was performed using GraphPad Prism (v11.0.2).

## Results

### Downregulation of SETD3 increases epithelial permeability and impairs junctional reassembly in model IEC monolayers

The functional roles of SETD3 in regulating junctional assembly and barrier properties in model IEC monolayers were investigated by performing siRNA-mediated knockdown of this methyltransferase in DLD-1 and HCA-7 human colonic epithelial cells. DLD-1 and HCA-7 are well-polarized epithelial cell lines that rapidly assemble adherens and tight junctions and are good models to study intestinal epithelial barrier regulation *in vitro* ^67–70^.

IEC transfection with SETD3 siRNA resulted in significant (60-90%) reduction of protein expression in both cell lines on day 4 post transfection (Figure 1 A, D). Interestingly, SETD3 depletion significantly decreased the amount of His73 methylated actin (Figure 1 A, E) indicating that this enzyme is essential for actin methylation in IECs. SETD3 siRNA knockdown did not cause significant decrease in viability of DLD-1 and HCA-7 cells (data not shown). The effect of SETD3 downregulation on IEC barrier integrity was determined by measuring the transepithelial electrical resistance (TEER) and transmonolayer flux of FITC-dextran (4 kDa). SETD3 knockdown significantly decreased TEER (Figure 1 B, E) and increased FITC-dextran permeability (Figure 1 C, F) on day 4 post-siRNA transfection in both DLD-1 and HCA-7 cell monolayers, suggesting increased permeability of the paracellular barrier. Next, we asked whether the observed weakening of the epithelial barrier is associated with the impaired junctional integrity in SETD3-depleted IECs. Immunofluorescent labeling and confocal microscopy were used to visualize junctional recruitment of TJ (ZO-1, claudin-7, claudin-4) and AJ (E-cadherin) proteins. In control DLD-1 and HCA-7 cell monolayers, all examined TJ and AJ proteins presented in a ‘chicken wire’ labeling pattern, typical for well-assembled epithelial junctions (Figure 2, Suppl. Figure 1, arrows). This labeling pattern was preserved in SETD3 depleted cells (Figure 2, Suppl. Figure 1, arrowheads). In agreement, immunoblotting analysis did not indicate changes in the total expression of different AJ and TJ proteins in SETD3-depleted DLD-1 and HCA-7 cells (Suppl. Figure 2). However, loss of SETD3 induced noticeable changes in IEC cell morphology, which manifested as a significant increase in epithelial cell size (Suppl. Figure 3A, arrows B) alongside a concomitant decrease in monolayer height (Suppl. Figure 3C, arrows D).

**Figure 1.**
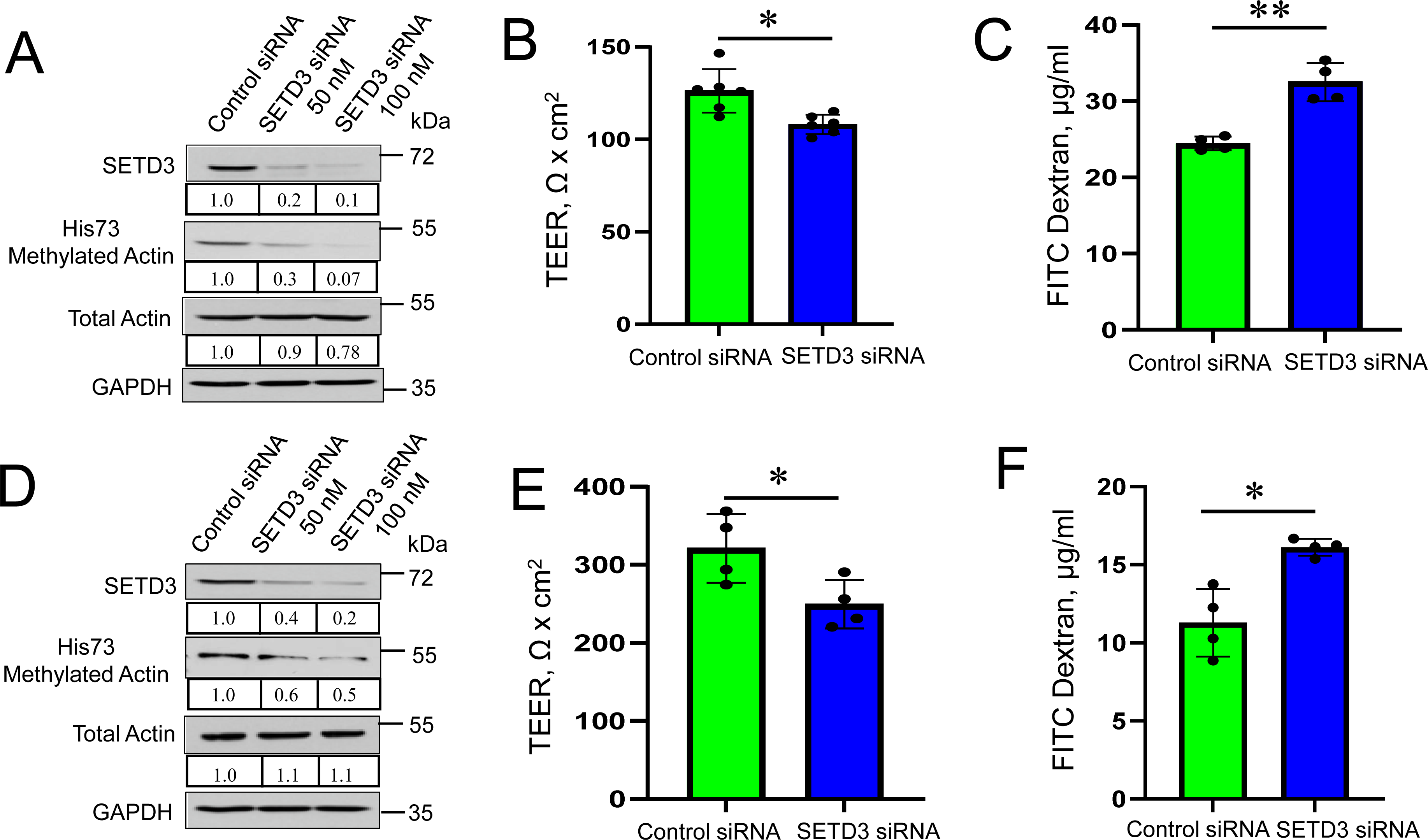
Downregulation of SETD3 increases paracellular permeability of model intestinal epithelial cell monolayers. (**A**,**D**) Immunoblotting analysis of SETD3 and methylated actin levels in DLD-1 (**A**) and HCA-7 (**D**) cells with siRNA-mediated knockdown of SETD3. (**B,E**) Transepithelial electrical resistance (TEER) of control and SETD3-depleted DLD-1 (**B**) and HCA-7 (**E**) cell monolayers on day 4 after siRNA transfection. (**C,F**) Trans-monolayer flux of the 4 kDa FITC-dextran in control and SETD3-depleted DLD-1 (**C**) and HCA-7 (**F**) cells on day 4 post siRNA transfection. Data are presented as means ± SE (n=4-6). *p< 0.05, **p< 0.01, as compared to control siRNA transfected cells.

**Figure 2.**
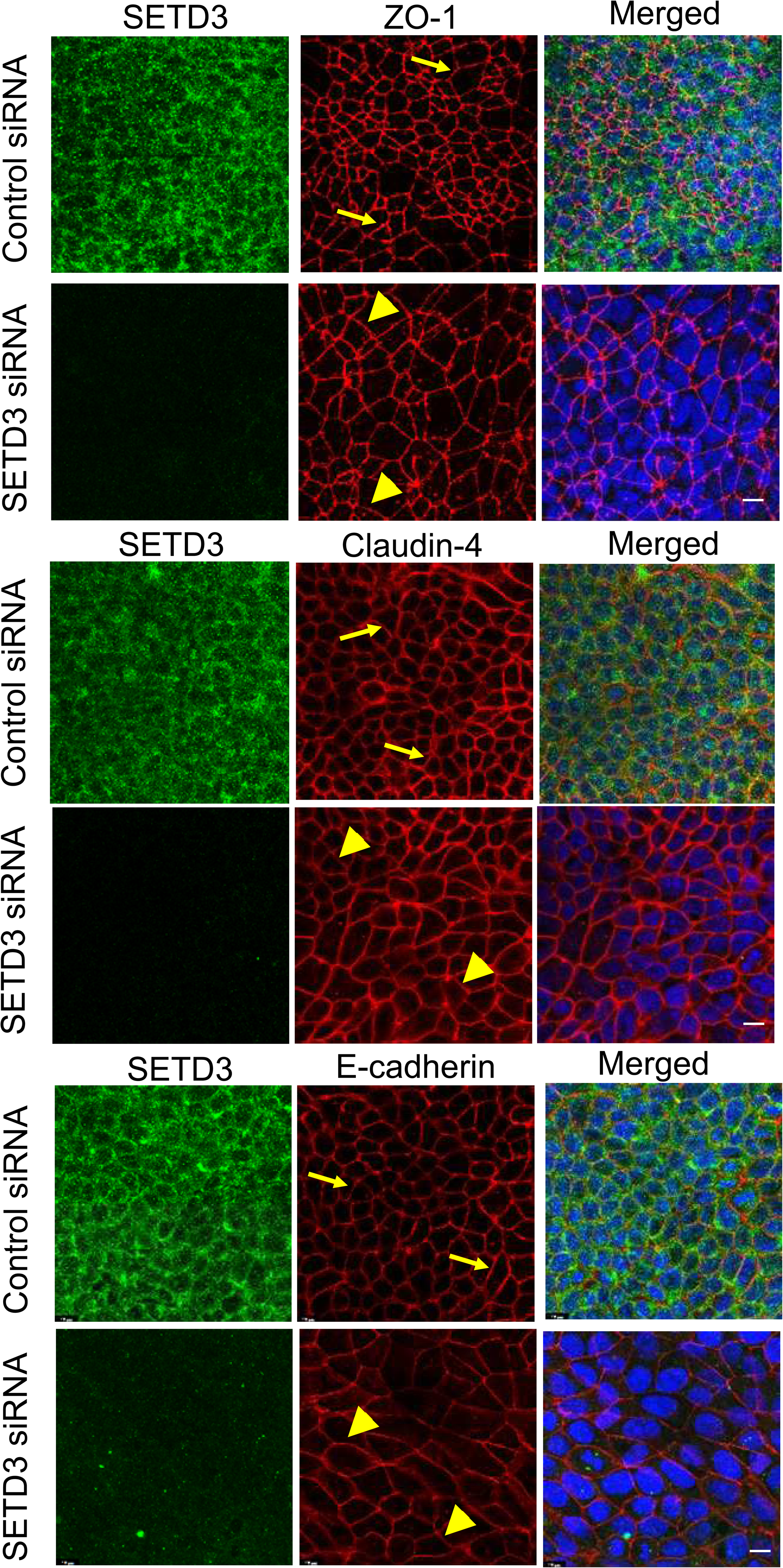
Depletion of SETD3 does not affect a steady state integrity of IEC junctions. A dual immunofluorescence labeling and confocal microscopy images of SETD3 (green) and ZO-1, claudin-7 or E-cadherin (red) in control and SETD3-depleted DLD-1 cell monolayers on day 4 after siRNA transfection. Nuclei are labeled with DAPI (blue). Arrows and arrowheads indicate intact TJ and AJ in control and SETD3-depleted cell monolayers, respectively. Images shown are representative of three independent experiments with multiple images taken per slide. Scale bars, 10 μm.

The intestinal epithelial layer is known to be a dynamic structure, characterized by robust remodeling of apical junctions under homeostatic conditions, and during epithelial injury and repair. As such, we sought to investigate the effects of SETD3 depletion on TJ/AJ reassembly using an extracellular calcium switch assay, in order to specifically study orchestrated junctional reassembly separate from other cellular processes such as cell division and migration ^33, 34, 36^. Confluent control and SETD3-depleted DLD-1 and HCA-7 cell monolayers were subjected to overnight extracellular calcium depletion to induce disassembly of cell-cell contacts, followed by returning the cells into the normal calcium medium to induce reassembly of apical junctions. Control DLD-1 and HCA-7 cells rapidly reestablished both TJ and AJ structures within 2 and 4 hours of extracellular calcium restoration, respectively (Figure 3, arrows). In contrast, TJ reassembly was significantly attenuated in SETD3-depleted DLD-1 (Figure 3A arrowheads, B) and HCA-7 (Figure 3C arrowheads, D) cells. Together, these findings suggest that SETD3 promotes reassembly of tight junctions and enhances epithelial barrier function in model IEC monolayers.

**Figure 3.**
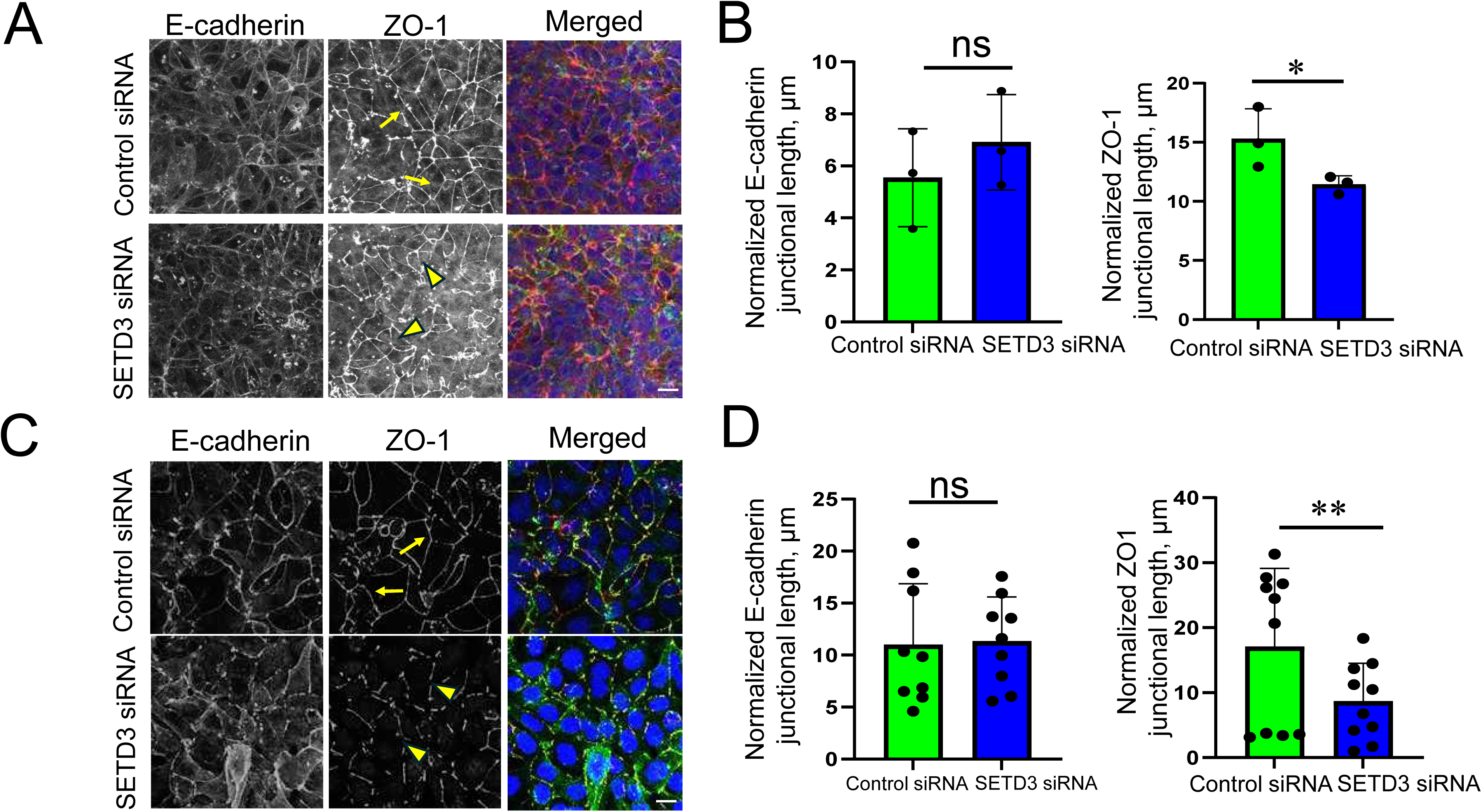
Depletion of SETD3 attenuates AJ and TJ reassembly during the extracellular calcium switch. Immunolabeling of E-cadherin (green) and ZO-1 (red) in control and SETD3-depleted DLD-1 (**A, B**) and HCA-7 (**C, D)** cells after 2 (DLD1) and 4 (HCA7) hours of extracellular calcium repletion, respectively. Representative confocal microscopy images (**A, C**) and quantification of the normalized junctional length (**B, D**) are shown. Arrows indicate AJ/TJ reassembly in control cells and arrowheads point at poorly assembled apical junctions in SETD3-depleted cells. Means ± SE (n=5). *p< 0.05, **p< 0.01, NS-non-significant. Scale bars, 10 μm.

### Loss of SETD3 impairs the integrity and reassembly of the perijunctional actomyosin belt

The dramatic alterations of the epithelial cell shape caused by SETD3 depletion strongly suggest the redistribution of cytoskeleton-dependent mechanical forces in cell monolayers. Therefore, we sought to determine whether loss of SETD3 affects the assembly of the epithelial actomyosin cytoskeleton. Cytoskeletal integrity was assessed by visualizing junction-associated F-actin, cytoskeletal motor NM IIA and the activated NM II pool associated with Thr18/Ser19 di-phosphorylated myosin light chains (ppMLC). Control DLD-1 cell monolayers contained a prominent perijunctional F-actin belt that was enriched in both NM IIA and ppMLC (Figure 4A, arrows). However, the staining intensity of F-actin, NM IIA and pp-MLC labeling in this circumferential belt was significantly decreased following SETD3 depletion (Figure 4A, arrowheads, B), indicating diminished assembly of the perijunctional actomyosin cytoskeleton. Importantly, total levels of NM IIA, NM IIC and ppMLC did not differ significantly between control and SETD3-depleted DLD-1 cells (Figure 4C). We complemented these data by performing live cell imaging of the perijunctional actin cytoskeleton by using a cell permeable F-actin probe, SPY650-FastAct-X ^71, 72^. In confluent control DLD-1 and HCA-7 cells, this probe predominantly labeled F-actin bundles at epithelial junctions (Suppl. Figure 4A,D). Incorporation of the SPY650-FastAct-X into the perijunctional F-actin was significantly lower in SETD3-depleted DLD-1 and HCA-7 cells (Suppl. Figure 4A,B,D,E). The overall intensity of the probe decreased with time, reflecting the dynamic remodeling of labeled actin filaments ^71, 72^, however, SETD3 depletion did not have consistent effect on the rate of the decrease of SPY650-FastAct-X intensity in DLD-1 and HCA-7 cell monolayers (Suppl. Figure 4C,F; Suppl. Movies 1-4).

**Figure 4.**
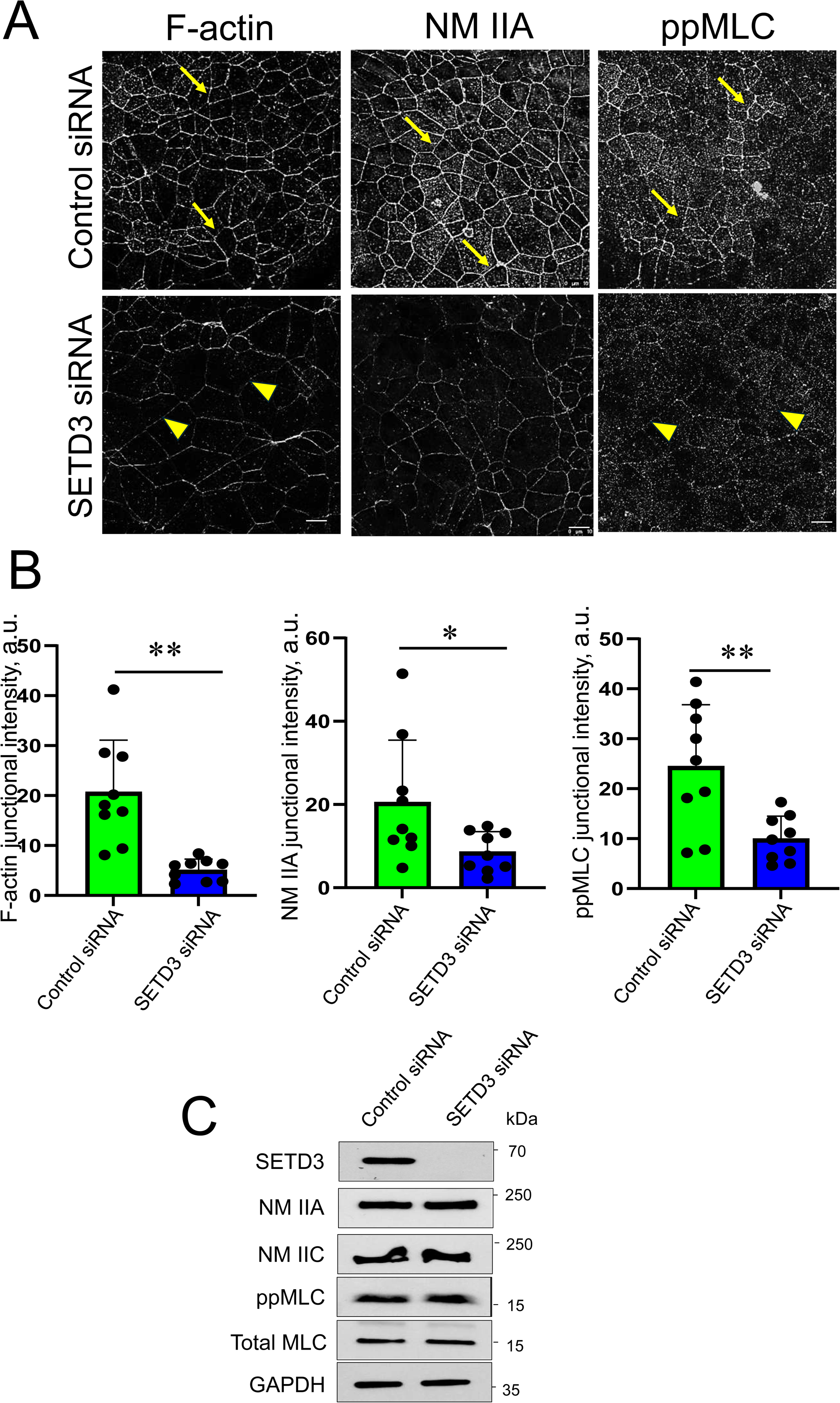
Loss of SETD3 impairs assembly of the perijunctional actomyosin belt. (**A, B**) Fluorescence labeling and confocal microscopy images of control and SETD3-depleted DLD-1 cells labeled for F-actin, NM IIA and ppMLC. (**A**) Representative images of three independent experiments with multiple images taken per slide. Arrows indicate perijunctional actomyosin structures in control epithelial cell monolayers. Arrowheads point at the decreased intensity of perijunctional actomyosin bundles in SETD3-depleted cells. (**B**) Quantification of signal intensities at apical junctions. Means ± SE (n=3-6). \*\**p* < 0.01, *p< 0.05. Scale bars, 10 μm. (**C**) Immunoblotting analysis of total expression of NM II motors and myosin regulatory light chains in cell lysates of control and SETD3-depleted DLD-1 cells.

In addition to examining the steady-state integrity of the perijunctional actomyosin belt, we investigated reassembly of this structure in control and SETD3-depleted cells during the calcium switch. Control DLD-1 cells efficiently restored the perijunctional actomyosin bundles after 2 hours of calcium repletion (Suppl. Figure 5A, arrows). By contrast, reassembly of the perijunctional actomyosin bundles was significantly delayed in SETD3-depleted cells (Suppl. Figure 5A, arrowheads, B). Together, our data suggests that increased permeability and altered cell morphology in SETD3 depleted intestinal epithelial cell monolayers is associated with abnormal assembly of the perijunctional actomyosin cytoskeleton.

### Loss of SETD3 increases intestinal permeability and impairs assembly of the epithelial actin cytoskeleton in vivo

Next, we sought to determine the relevance of our findings *in vivo* and characterize the gut barrier integrity and intestinal epithelial junction-associated cytoskeleton in mice with total knockout of SETD3 (SETD3 KO). As expected, SETD3 KO mice demonstrated a complete loss of SETD3 protein expression in the colonic epithelium, which was accompanied by a complete loss of His73 actin methylation (Figure 5A). This data highlights SETD3 as the major actin methyltransferase in the mouse colonic epithelium. We also examined the architecture of the IEC cytoskeleton by labeling colonic sections with antibodies selectively recognizing β-cytoplasmic and γ-cytoplasmic actins ^26^. Both cytoplasmic actins were abundant at the apical pole of colonic IECs (Figure 5B, C, arrows). Interestingly, loss of SETD3 selectively decreases the assembly of apical β-cytoplasmic actin without having an effect on the other actin paralog (Figure 5B, C, arrowheads). SETD3 KO mice appeared healthy and did not show any signs of gastrointestinal pathology (data not shown). Furthermore, SETD3 knockout did not affect general mucosal architecture in proximal and distal colon according to Hematoxylin & Eosin (H&E) staining (Suppl. Figure 6, arrows). Yet, SETD3 KO animals were characterized by a significant increase in gut barrier permeability as compared to their wild-type littermates (Figure 6A). The increased intestinal permeability of SETD3 KO mice was not associated with significant alterations of the tight junction integrity in the colonic epithelium as indicated by immunofluorescence labeling of different TJ proteins (Figure 6B, arrows).

**Figure 5.**
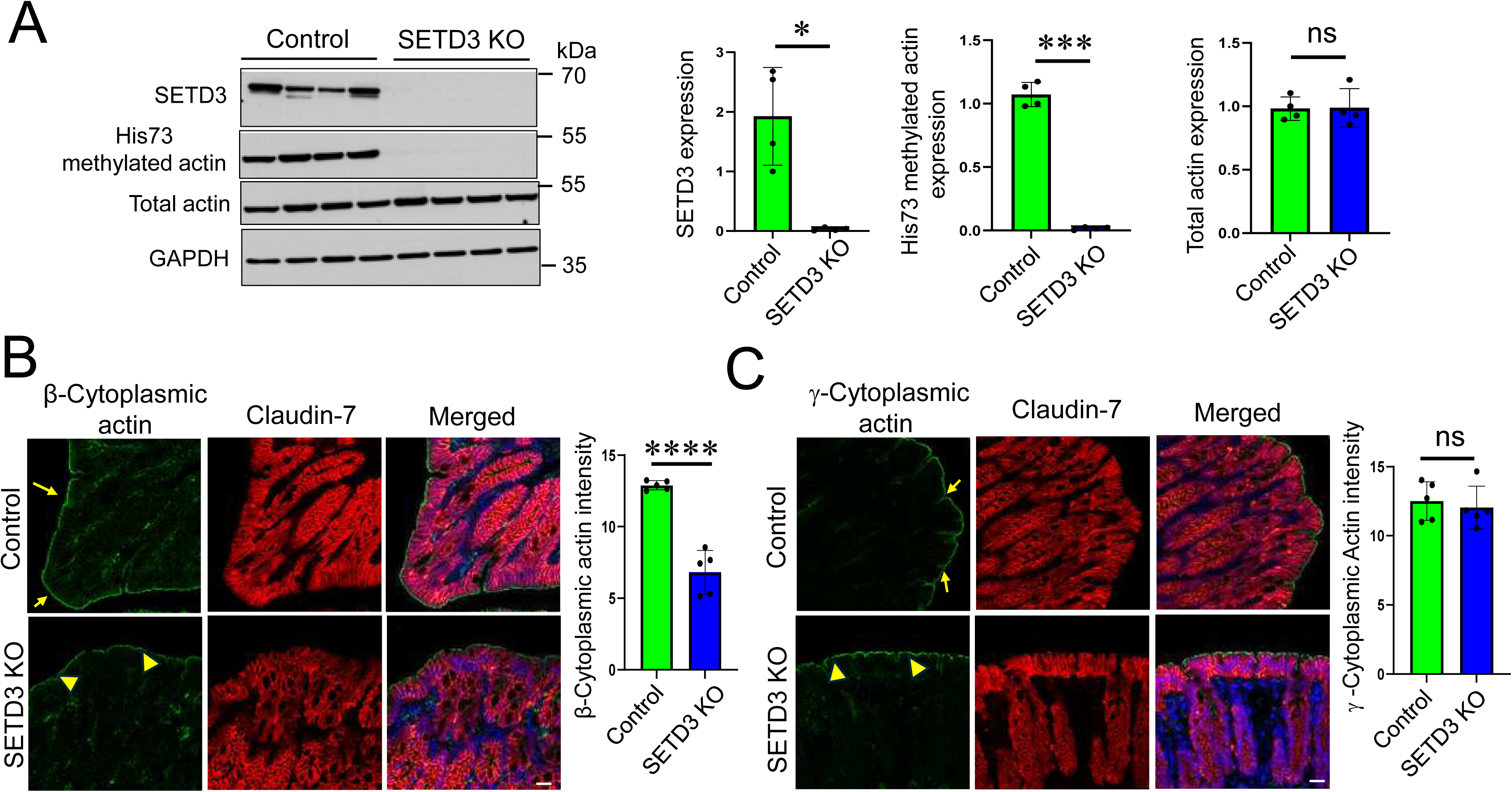
Loss of SETD3 in mice downregulates the level of His73 methylated actin and decreases assembly of β cytoplasmic actin cytoskeleton in colonic mucosa. (**A**) Immunoblotting analysis of SETD3 and His73-methylated actin expression in colonic epithelial scrapes of control and SETD3 null mice. Original immunoblot images and densitometric quantification are shown. Means ± SE (n=4). *p< 0.05, ***p< 0.001, ns, not significant. (**B, C**) Normal colonic sections of control and SETD3 KO mice were dual-immunolabeling for either β-cytoplasmic actin (green, **B**), γ-cytoplasmic actin (green, **C**) and epithelial cell marker, claudin-7 (red). Arrows show assembly of the apical epithelial actin cytoskeleton in control mice. Arrowheads highlight the diminished assembly of β-cytoplasmic actin-based apical cytoskeleton in colonic epithelium of SETD3 KO mice. Means ± SE (n=5). ****p< 0.0001, ns, not significant. Scale bars, 25 μm.

**Figure 6.**
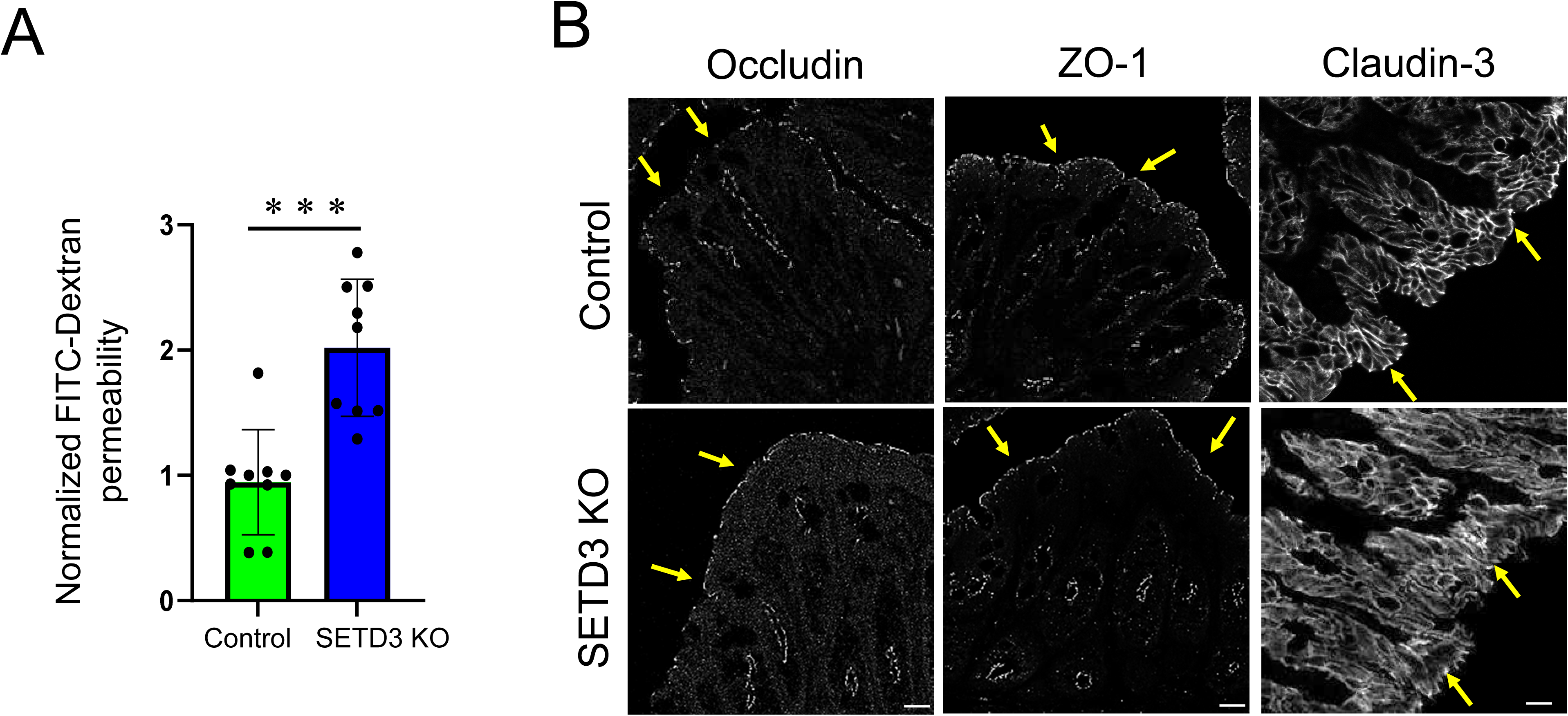
SETD3 knockout mice have increased intestinal permeability without marked alterations in the assembly of tight junctions. (**A**) Gut to blood passage of 4 kDa FITC-dextran in control and SETD3 KO mice. Means ± SE (n=9). ***p< 0.001. (**B**) Immunolabeling of occludin, ZO-1, and claudin-3 in the colonic mucosa of control and SETD3 KO mice. Arrows show normal TJ assembly in colonic epithelium control and SETD3 knockout animals. Scale bar, 25 μm.

Next, we sought to determine whether loss of SETD3 *in vivo* affects normal IEC homeostasis, by examining cell proliferation and abundancy of different IEC lineages. Immunolabeling with the Ki-67 cell proliferation marker revealed significantly increased numbers of Ki-67 positive cells in the colonic crypts of SETD3 KO mice (Figure 7A). Likewise, loss of SETD3 resulted in the increased expression of colonic stem cell marker, OLFM4 (Figure 7B). To complement this data, we compared abundance of different secretory epithelial cell lineages in the intestinal mucosa of SETD3 KO and control mice by immunolabeling with different secretory lineage markers. No significant differences were observed in Goblet, Tuft, and Paneth cell numbers between SETD3 KO and control mice based on immunolabeling of MUC2, DCLK1 and lysozyme, respectively (Suppl. Figure 7). Together, this data indicates that loss of SETD3 impairs gut barrier integrity and assembly of the epithelial actin cytoskeleton in normal intestinal mucosa, alongside a concomitant increase in IEC stemness and proliferation.

**Figure 7:**
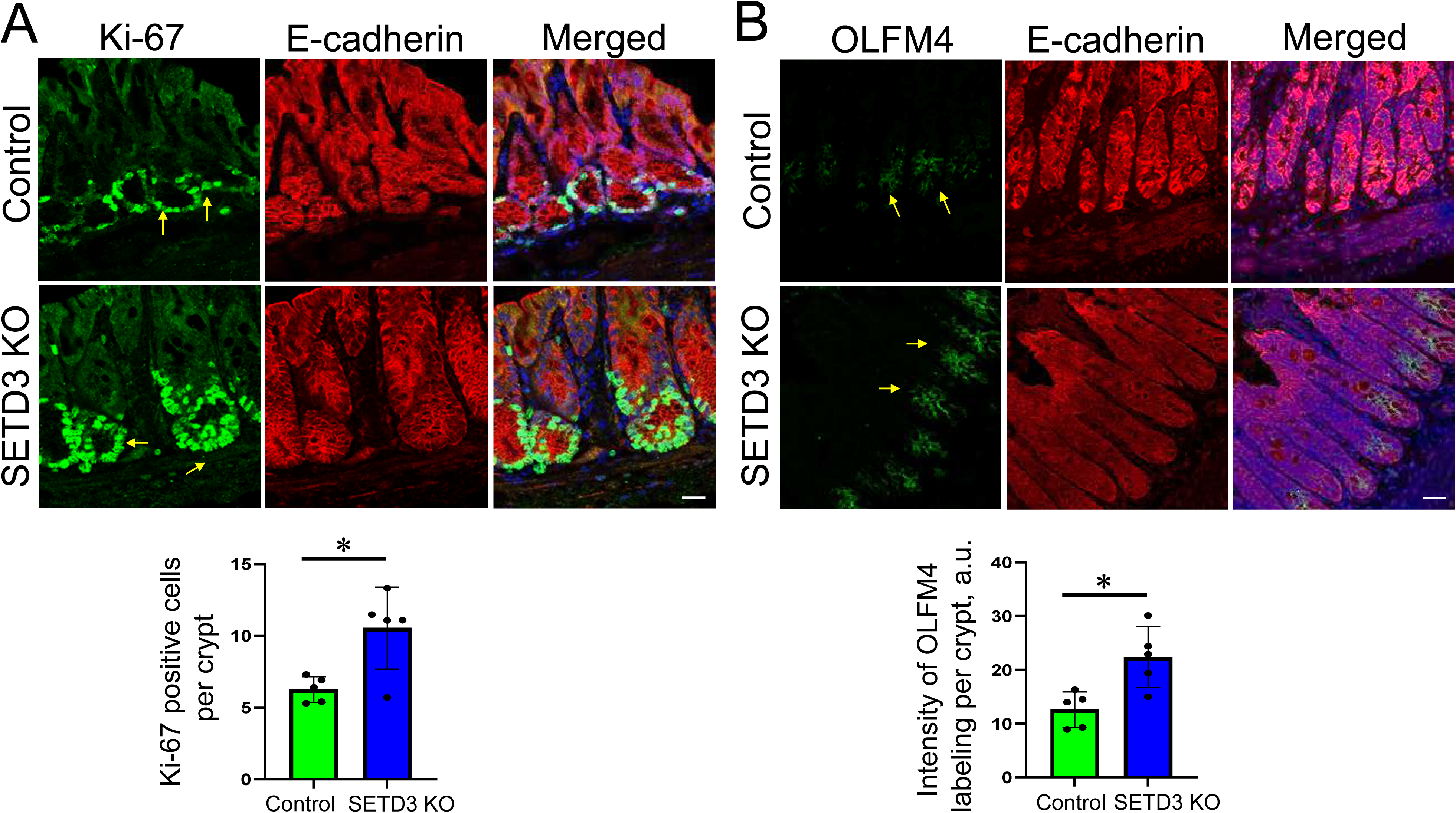
Los of SETD3 increases colonic epithelial cell proliferation and stemness *in vivo*. **(A)** Dual immunolabeling of Ki-67 (green) and E-cadherin (red) and (**B**) dual immunofluorescence labeling of OLMF4 (green) and E-cadherin (red) in normal colonic sections of control and SETD3 knockout mice. Representative images and image quantification are shown. Arrows point to Ki-67 and OLMF4 positive cells Means ± SE (n=5). *p< 0.05. Scale bars, 25 µm.

### SETD3 knockout alters the transcription program in intestinal epithelium in vivo

Given our findings of a leaky gut barrier and enhanced IEC proliferation in SETD3 KO mice, we sought to determine if these homeostatic changes are linked to changes in the transcriptional program of IECs. Bulk RNAseq analysis was performed using ileal and colonic epithelial cells isolated from SETD3 KO mice and their wild type littermates. While, a canonical correspondence analysis, which clusters individual samples based on their gene expression profiles, revealed partial overlap of transcriptional landscapes, (Suppl. Figure 8A), there are significant changes in transcriptional patterns between wild type ileal and colonic sections and their SETD3 knock out counterparts. The Permanova analysis results in R2=0.19 and p=0.021 for ileal cells and R2=0.26, p=0.001 for colonic cell transcriptomes, respectively. Among these observed changes, loss of SETD3 resulted in changes in downregulation of 128 genes and upregulation of 25 genes in colonic epithelium, along with downregulation of 156 and upregulation of 22 genes in ileal epithelium (Suppl. Figure 8B,C; Supplementary Table 1). In particular, pathway enrichment analysis indicated that transcripts related to responses to chemokines and lymphocyte chemotaxis were upregulated in colonic mucosa of SETD3 KO mice (Suppl. Figure 8D). Additionally, genes related to immune responses, secretory granules and host defense to bacteria were upregulated, while genes related to protein methylation and integrin-based adhesion were downregulated in SETD3 knockout ileal epithelium (Suppl. Figure 8E).

### Loss of SETD3 increases sensitivity to DSS colitis in mice

Given our finding of the upregulation of immune response related genes in SETD3 deficient IECs, we next asked if this methyltransferase regulates intestinal inflammation *in vivo*. Intestinal inflammation was induced in SETD3 KO mice and their wild type littermates via the administration of 2.5% dextran sodium sulfate (DSS) in drinking water. SETD3 KO mice displayed more severe disease-like symptoms as compared to control mice, which manifested as greater body weight loss (Figure 8A) and higher disease activity index (Figure 8B). H&E staining revealed marked signs of mucosal injury and inflammation including epithelial ulceration, edema and leukocyte infiltration caused by DSS colitis (Figure 8C). Such tissue injury was significantly higher in the middle colon of SETD3 mice as compared to their wild type littermates (Figure 8D).

**Figure 8:**
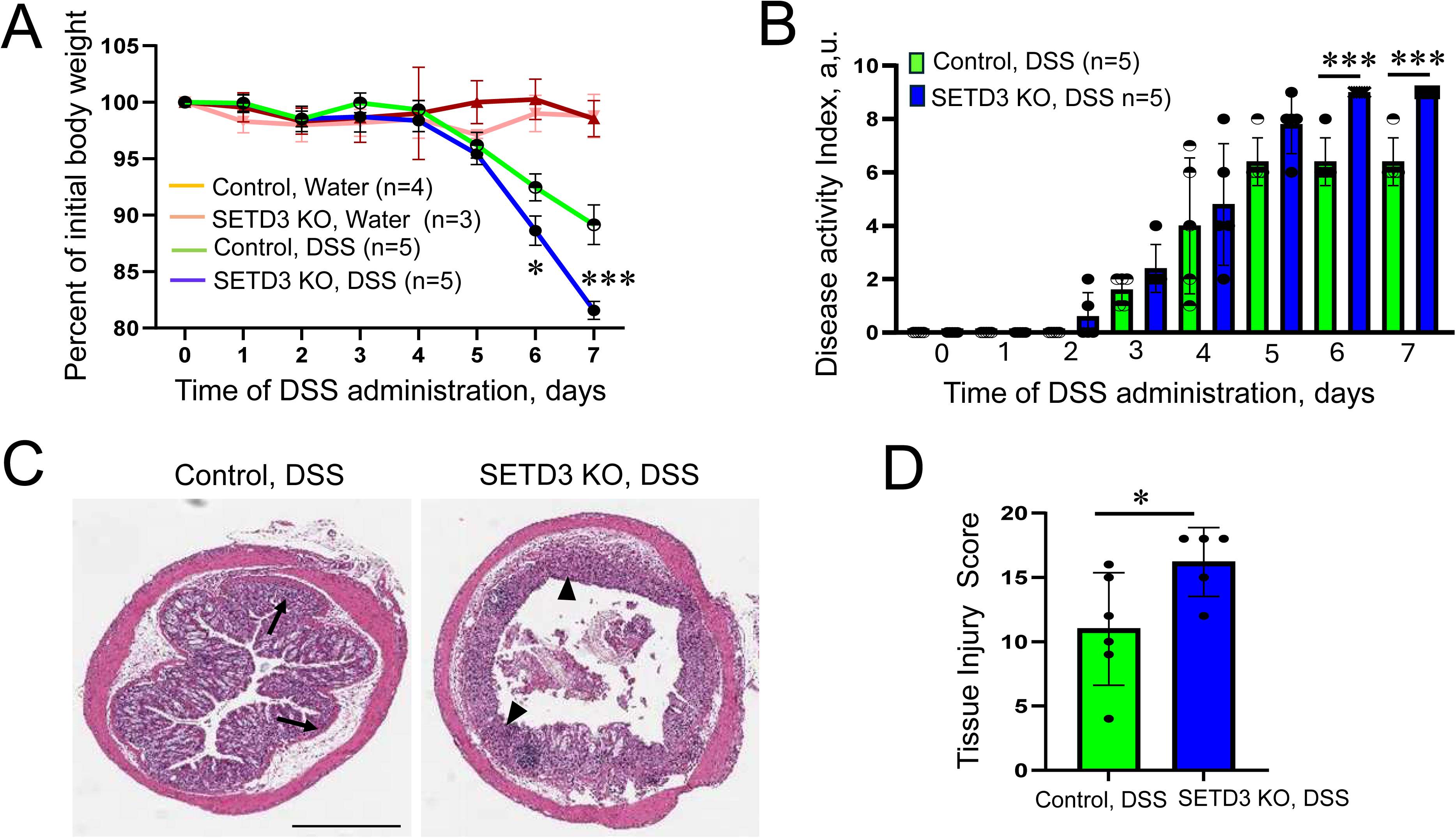
Knockout of SETD3 exacerbates disease severity during DSS-induced colitis. Control and SETD3 KO mice were exposed to 2.5% DSS in drinking water, or regular drinking water as a control. (**A**) Body weight loss and (**B**) the disease activity index were calculated for the duration of DSS treatment. (**C**) Hematoxylin & eosin-stained middle colonic sections of DSS treated control and SETD3 KO mice. Arrows show leukocyte infiltration and edema in DSS-treated animals. Arrowheads highlight epithelial erosion in SETD3 KO mice. (**D**) The calculated tissue injury score for the colonic section of DSS-exposed animals. Mean ± SE, (n = 3-6). \**p* < 0.05, \*\*\**p* < 0.0001. Scale bar, 900 μm.

### SETD3 KO mice show higher inflammatory cell infiltration in the colonic mucosa during DSS-induced colitis

Next, we sought to have a deeper characterization of intestinal inflammation in SETD3 null and control mice by examining immune cell infiltration in the colonic tissue during DSS colitis. Specifically, we examined the abundance of T-cells, macrophages and neutrophils by immunofluorescence labeling for their specific markers, CD3, F4/80 and Ly6G, respectively. We did not find differences in immune cell populations in the normal colon of SETD3 KO and control mice, indicating that loss of SETD3 does not cause spontaneous intestinal inflammation (Figure 9 A,B). As expected, DSS colitis resulted in an enhancement of all immune cells examined (Figure 9 A,B). The numbers of T-cells, macrophages and neutrophils were significantly higher in the colonic mucosa of DSS-treated SETD3 KO mice as compared to their control littermates (Figure 9 A,B). Thes data

**Figure 9:**
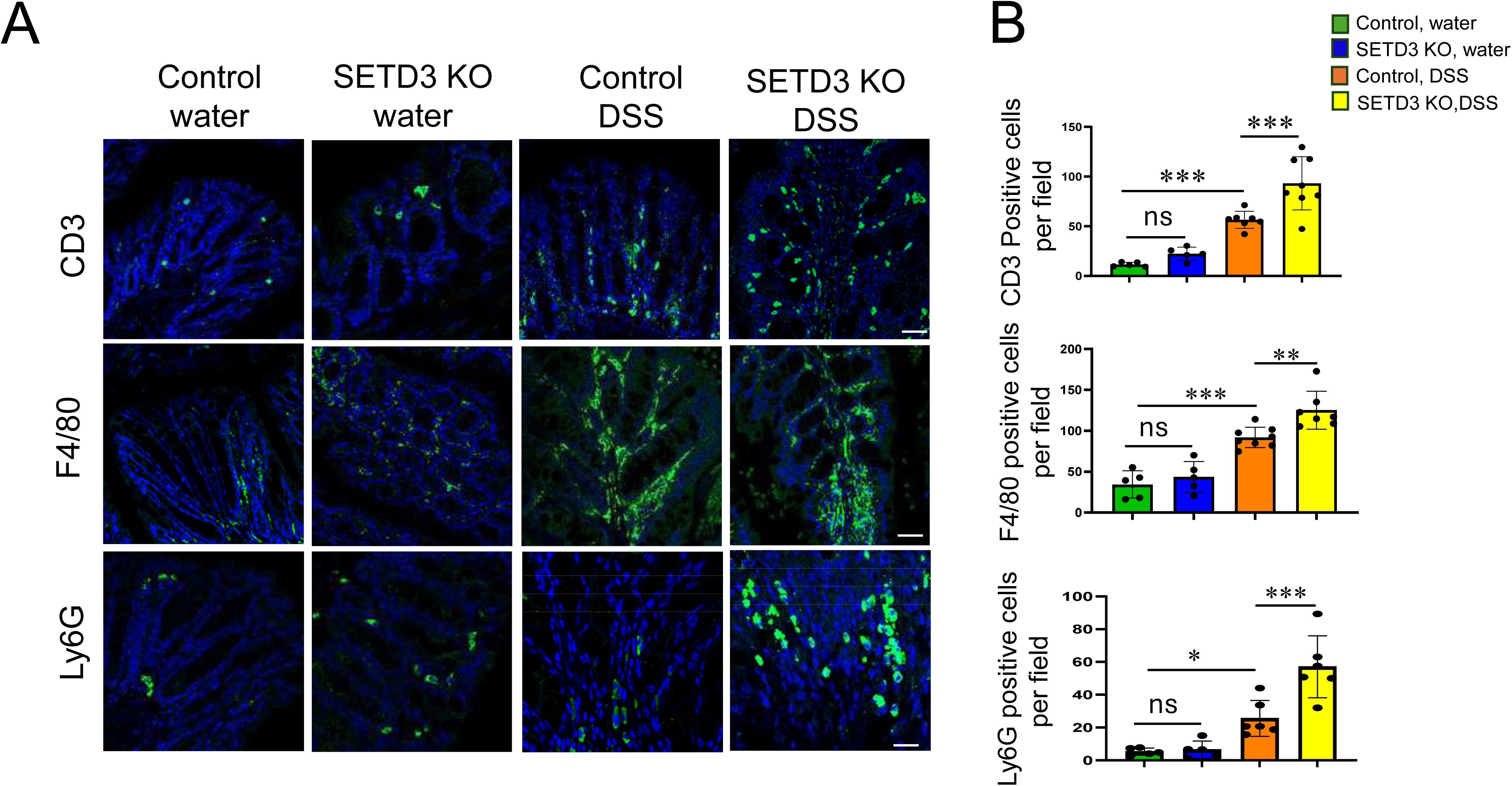
Loss of SETD3 exaggerates leukocyte infiltration in the colonic mucosa during DSS colitis. Control and SETD3 KO mice were exposed to 2.5 % DSS in drinking water, or regular water for 7 days. Harvested colonic samples were immunofluorescence labeled for the specific markers of different immune cell types (green): CD3 (T cells), F4/80 (macrophages), and Ly6G (neutrophils). All tissue sections were counter-labeled with nuclear stain, DAPI (blue). Representative images (A) and quantification of the immune cell numbers (B) are shown. Mean ± SE (n = 5), \**p* < 0.05, \*\**p* < 0.01, \*\*\**p* < 0.005. Scale bar, 25 μm.

Together, these data indicate that loss of SETD3 exaggerates intestinal epithelial damage, and mucosal inflammation during experimental colitis.

## Discussion

His73 methylation is a common posttranslational modification of mammalian actin molecules that modulates different aspects of actin plasticity such as ATP hydrolysis, phosphate release and actin filament polymerization ^46, 47, 73^. Emerging evidence indicates that this PTM is involved in the regulation of different cellular processes in normal and cancer cells ^74–76^. The present study identifies the SETD3 methyltransferase responsible for His73 actin methylation as a novel regulator of the intestinal epithelial barrier and a potent suppressor of intestinal inflammation.

The barrier protective function of SETD3 is supported by our studies of paracellular permeability in model IEC monolayers and in mice, which consistently indicates that barrier integrity is compromised by SETD3 depletion (Figures 1 & 6A). SETD3 controls permeability of IEC monolayers to ions and larger molecules, based on TEER and trans monolayer FITC-dextran flux measurements (Figure 1). Interestingly, the increased epithelial permeability caused by loss of SETD3 was not due to major alterations in the structure and molecular composition of TJs and AJs (Figures 2 & 6B, Suppl. Figures 1 & 2). The underlying mechanism involved a defective assembly of the prejunctional actomyosin cytoskeleton. This conclusion is supported by several lines of evidence. First, is the decreased intensity of perijunctional actin filaments in SETD3-deficient confluent IEC monolayers and mouse colonic mucosa (Figures 4 & 5; Suppl. Figure 4). Importantly that such diminished assembly of the perijunctional F-actin belt was observed in live IECs and well as in fixed epithelial cells and colonic tissues. Second is decreased recruitment and activation of the NM IIA motor at the perijunctional actomyosin belt (Figure 4). Third, is vivid alterations of cell shape (expansion of the apical surface, and reduction of the cellular height), which is indicative of the altered distribution of actomyosin forces (Suppl. Figure 3). While no previous studies investigated the effects of SETD3 depletion on the integrity of junction-associated actin filaments, our findings are consistent with previously reported diminished density the actin filament network observed in SETD3-deficient human HAP1 cells ^45^ and C2C12 mouse myoblasts ^77^, as well as decreased F-actin intensity in mouse cortical neurons ^52^. Importantly, loss of SETD3 not only decreased robustness of the perijunctional actomyosin cytoskeleton in differentiated IEC monolayers but also attenuated the reformation of the junction-associated cytoskeleton during extracellular calcium switch (Suppl. Figure 5). Such delayed cytoskeletal reassembly resulted in the delayed reformation of TJs but not AJs (Figure 3), thereby highlighting the roles of SETD3 in regulating junctional remodeling.

While actin-dependent regulation of epithelial junctional integrity is very robust *in vitro*, such regulation could be less obvious *in vivo* where IECs experience lower mechanical forces from softer tissues ^26, 39^. It is important, therefore, to observe alterations of gut barrier integrity and IEC homeostasis in SETD3 null mice. Specifically, loss of SETD3 increases permeability of the normal gut barrier in mice in parallel to decreased assembly of the apical actin cytoskeleton (Figures 5 & 6). Additionally, SETD3 KO mice show increased proliferation of colonic epithelial cells (Figure 7) yet is not accompanied by decreased differentiation of the major secretory epithelial cell types (Suppl. Figure 7). This data contrasts with previous studies, which highlight SETD3 as an activator of endoderm differentiation in mouse embryonic stem cells ^78^ as well as a regulator of muscle cell differentiation ^49, 79^. Therefore, SETD3 may be essential for cell differentiation in a tissue-specific fashion. What are the possible reasons for the mild phenotype of SETD3 knockout in the gut? It is not because of functional redundancy with other methyltransferases, since SETD3 appears to be the major, if not sole mediator, of His73 actin methylation in the intestinal epithelium (Figure 5). Instead, it could be that the functional consequences behind the loss of H73 methylation do not prevent actin filament assembly altogether but rather change the dynamics of this process. It is likely that cells and organisms can adapt to modest alterations in actin filament dynamics.

Transcriptomic analysis of SETD3-deficient IECs suggests that SETD3 may play a role in regulating gene expression in the intestinal epithelium (Suppl. Figure 8). This is consistent with previously reported transcriptional alterations observed in SETD3 depleted mouse embryonic stem cells ^78^, mouse myoblasts ^49^, T cells ^80^ and breast cancer cell lines ^51^. This data raises an important question about the mechanisms of SETD3-regulated gene expression. Generally, two mechanisms could be envisioned. One involves actin methylation-dependent regulation of the transcription activity of nuclear actin. Indeed, β-actin methylation is an emerging regulator of gene expression that impacts chromatin remodeling, histone acetylation, and enhancer dependent gene transcription ^81, 82^. In fact, a recent study demonstrated that SETD3 dependent methylation controls β-actin interactions with different complexes involved with transcriptional regulation in colon cancer cells ^83^. An alternative mechanism involves SETD3-dependent modifications of non-actin targets. Emerging evidence suggests that in addition to actins, SETD3 may interact with various proteins including bromodomain containing protein 2 ^84^, mRNA splicing factors ^85^, and CDH1 DNA-binding protein ^86^. Future studies are required to delineate the exact mechanisms of SETD3 dependent regulation of the transcriptional program in IECs.

Increased animal sensitivity to acute DSS-induced colitis represents the most vivid consequence of SETD3 knockout *in vivo*. This involves exaggerated disease symptoms (Figure 8), increased accumulation of different immune cells in the colonic mucosa and accelerated IEC death (Figure 9). Interestingly, the response of SETD3 KO mice to DSS colitis recapitulates the increased colitis sensitivity observed in mice with intestinal epithelial specific deletion of β-actin ^26^. This may suggest the possibility of shared mechanisms between the protective roles of SETD3 and β-actin in the inflamed intestinal epithelium. Such mechanisms may involve stabilization of the intestinal epithelial barrier, as SETD3 KO mice and mice with β-actin depletion in the intestinal epithelium show exaggerated gut barrier permeability under homeostatic conditions.

In conclusion, the present study identifies a novel mechanism that regulates the integrity of the intestinal epithelial barrier and remodeling of apical junctions, while playing a protective role during acute intestinal inflammation. This mechanism involves SETD3-dependent regulation of the junction-associated actomyosin cytoskeleton. SETD3 appears to have multiple functions in the intestinal epithelium by mediating His73 methylation of actin, thereby controlling paracellular permeability of the intestinal epithelial barrier in vitro and in vivo, mediating assembly of junction-associated actomyosin cytoskeleton, and regulating the transcriptional program of mouse intestinal epithelial cells. Our study adds to the complexity of cytopskeletal mechanisms of epithelial barrier regulation by highlighting the importance of posttranslational modifications of actin molecules. It also identified a novel modulator of intestinal inflammation and injury that may also contribute to the pathogenesis of different inflammatory and immune disorders.

## Acknowledgments

The authors thank Drs. Or Gozani and Jan Carette for providing breeding pairs of SETD3 null mice and anti His73 methylated actin antibody. The authors also thank Dr. Fan Zhang for contributing to the initial stage of this study and Dr. Min Hui Lim, a Program manager for the Cleveland Clinic Research Genomic Core for facilitating RNA sequencing. We thank Marlo Nicolas for performing a pathology analysis and scoring of mouse tissue sections. This work was supported by National Institute of Health grant R01DK137822 to A.I.I. This work utilized the Leica SP8 confocal microscope in the Imaging Core of the Cleveland Clinic Research that was purchased with funding from National Institutes of Health SIG grant 1S10OD019972-01.

## Author Contributions

P.M. performed experiments, analyzed and interpreted data and prepared figures. N.N and S.L. performed experiments and participated in data analysis. A. Z. helped with live cell imaging and analyzed the imaging data. N.S. performed bulk RNA sequence analysis. A.D. participated in data analysis and experiments. A.I.I. conceived the study, supervised the project and wrote the manuscript. All authors have read and approved the manuscript.

## Supplementary Figures

**Supplementary Figure 1. Depletion of SETD3 does not affect a steady state AJ and TJ morphology in HCA-7 cell monolayers.**

Dual immunofluorescence labeling and confocal microscopy images of SETD3 (green) and ZO-1, claudin-4 or E-cadherin (red) in control and SETD3-depleted HCA-7 cell monolayers on day 4 after siRNA transfection. Nuclei are labeled with DAPI (blue). Arrows and arrowheads indicate intact TJ and AJ in control and SETD3-depleted cell monolayers, respectively. Scale bar, 10 μm.

**Supplementary Figure 2. Loss of SETD3 does not affect expression of different junctional proteins in model IEC monolayers.**

DLD-1 (**A**) and HCA-7 (**B**) cells were transfected with either control or SETD3-specific siRNAs and harvested to prepare total cell lysates on day 4 after transfection. Immunoblotting analysis shows expression of different AJ and TJ proteins in control and SETD3-depleted intestinal epithelial cells.

**Supplementary Figure 3. Depletion of SETD3 increases cell size and decreases cell height in DLD-1 cell monolayers.**

(**A**) Dual immunofluorescence labeling and confocal microscopy images of SETD3 (green) and ZO-1(red) in control and SETD3-depleted DLD1 cell monolayers on day 4 after siRNA transfection. Arrows show the increased size of SETD3 depleted cells. (**B**) Quantification of epithelial cell size. (**C**) xz sections of control and SETD3-depleted DLD-1 cell monolayers dual immunolabeled for E-cadherin (green) and claudin-7 (red). Nuclei are labeled with DAPI (blue). Arrow points at the decreased height of SETD3-depleted cell monolayer. (**D**) Quantification of epithelial cell height in control and SETD3-deficient DLD1 cell monolayers. Means ± SE (n=9). *p< 0.05, ***p< 0.005, as compared to control siRNA transfected cells. Scale bars, 25 μm (A), 5 μm (B)

**Supplementary Figure 4.** Live-cell imaging analysis of the actin cytoskeleton using a SPY650-FastAct-X probe. Control and SETD3-depleted DLD-1(**A-C**) and HCA-7 (**D-E**) cell monolayers were incubated with the SPY650-FastAct-X probe for 2 h. The unbound probe was washed out and the labeled cells were imaged every 1 min for 6 h. (**A,D**) Snapshots of initial probe labeling. (**B,E**) Average signal intensity of the SPY™650-FastAct_X probe over the 1h imaging period in DLD-1 (**B**) and HCA-7 (**E**) cells. (**C,F**) Normalized signal intensities of the SPY650-FastAct-X probe over time in DLD-1 (**C**) and HCA-7 (**F**) cells. Mean ± SE. ****p< 0.0001 (n=60).

**Supplementary Figure 5. Loss of SETD3 impairs assembly of the perijunctional actomyosin belt.**

Fluorescence labeling of F-actin (green) and NM IIA (red) in control and SETD3-depleted DLD-1 cells after 2 hours of extracellular calcium repletion. Representative confocal microscopy images (**A**) and quantification of the normalized junctional length **(B)** are shown. Arrows indicate AJ/TJ reassembly in control cells and arrowheads point at poorly assembled apical junctions in SETD3-depleted cells. Mean ± SE (n = 3), **p < 0.01, ***p < 0.001. Scale bar, 10 μm.

**Supplementary Figure 6. SETD3 knockout does not cause dramatic alterations in colonic mucosal architecture in mice.** Hematoxylin & eosin-stained distal & proximal colonic sections of control and SETD3 KO mice. Arrows point on the normal architecture of colonic mucosa in SETD3 null mice and their wild type controls. Scale bar, 300 μm.

**Supplementary Figure 7: SETD3 knockout in mice does not affect secretory epithelial lineages differentiation in the intestinal mucosa.**

Immunolabeling of a Goblet cell marker, MUC2 (**A,** green), and tufts cell marker, DCLK1 (**B**, green) in colonic sections, as well as immunolabeling of a Paneth cell marker, lysozyme (**C**, green), in ileal sections of control and SETD3 KO mice. All secretory cell markers are counterstained with general epithelial markers, E-cadherin and β-catenin (red) and nuclear DAPI (blue). Representative images and image quantification are shown. Means ± SE (n=5). ns, not significant. Scale bars, 25 μm.

**Supplementary Figure 8: Loss of SETD3 induces transcription reprogramming of intestinal epithelial cells in mice.** (**A**) Principal Component Analysis (PCA) of gene expression profiles in colonic and ileal epithelial cells isolated from SETD3 KO and wild type control mice. Volcano plots showing gene expression in (**B**) colonic and (**C**) ileal epithelium of SETD3 KO as compared to control mice. Significant differentially (p-value and log2FC cut-off) expressed genes are highlighted by red inverted triangles. (**D**, **E**) Bubble plots representing pathway enrichment analysis of genes with statistically significant differences in expression between SETD3 KO and control mice in the (**D**) colonic and (**E**) ileal epithelium. Each bubble corresponds to a specific pathway, with size indicating the gene ratio and color representing the significance of the enrichment. The analysis was performed using isolated IECs from 5 mice/group.

## List of Abbreviations

AJs: adherens junctions
FITC: fluorescein isothiocyanate
KO: knockout
MLC: myosin light chain
NM II: non-muscle myosin II
PTMs: posttranslational modifications
SETD3: SET domain protein 3
TEER: transepithelial electrical resistance
TJs: tight junctions
ZO: zonula occludens

