## Supplemental figure1 for "SETD3 ACTIN HISTIDINE METHYLTRANSFERASE REGULATES INTESTINAL EPITHELIAL BARRIER INTEGRITY AND MUCOSAL INFLAMMATION"

### Slide 1
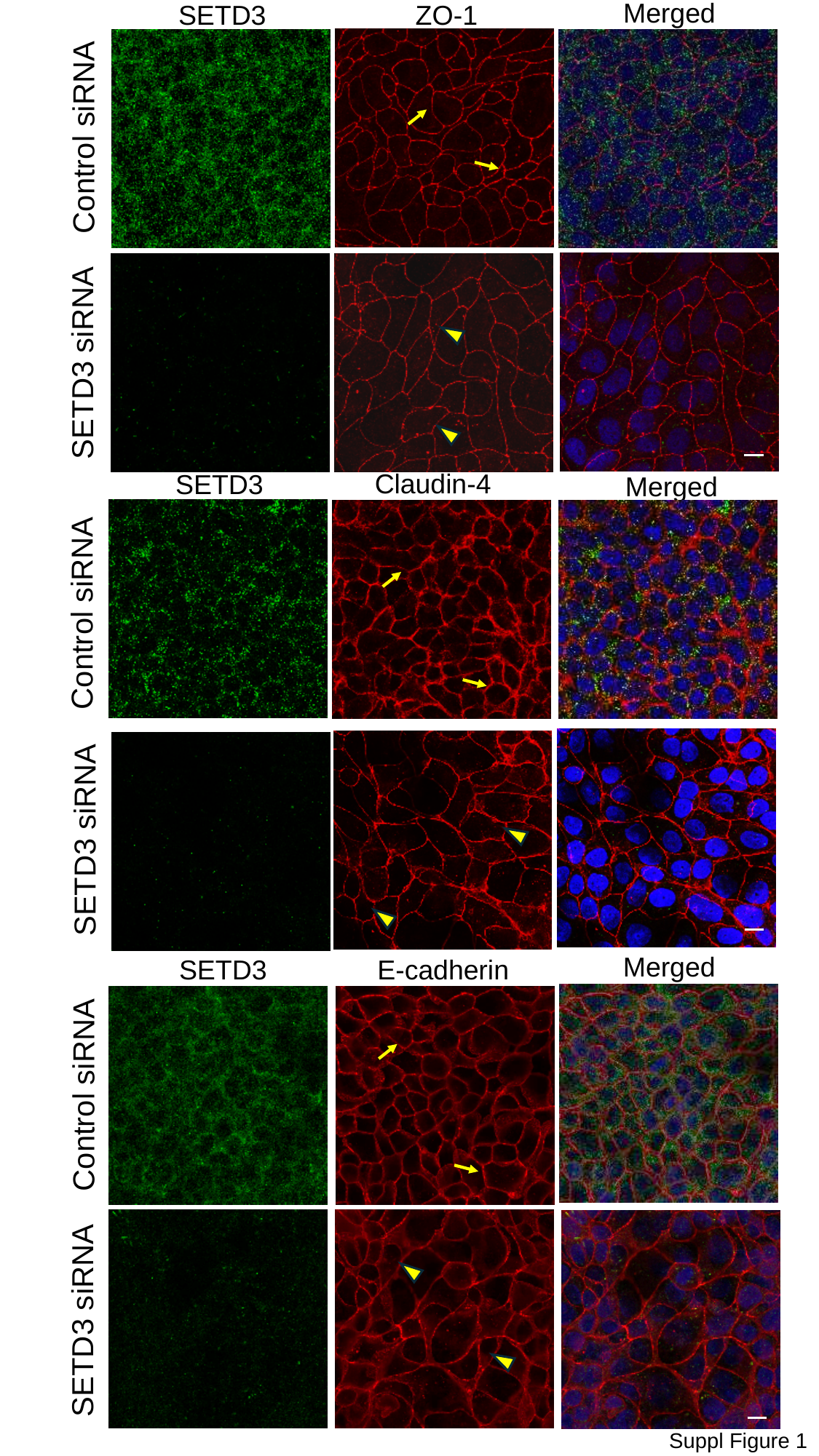

Merged
SETD3
ZO-1
Control siRNA
SETD3 siRNA
Claudin-4
SETD3
Merged
Control siRNA
SETD3 siRNA
Merged
E-cadherin
SETD3
Control siRNA
SETD3 siRNA
Suppl Figure 1
