## Supplemental figure2 for "SETD3 ACTIN HISTIDINE METHYLTRANSFERASE REGULATES INTESTINAL EPITHELIAL BARRIER INTEGRITY AND MUCOSAL INFLAMMATION"

### Slide 1
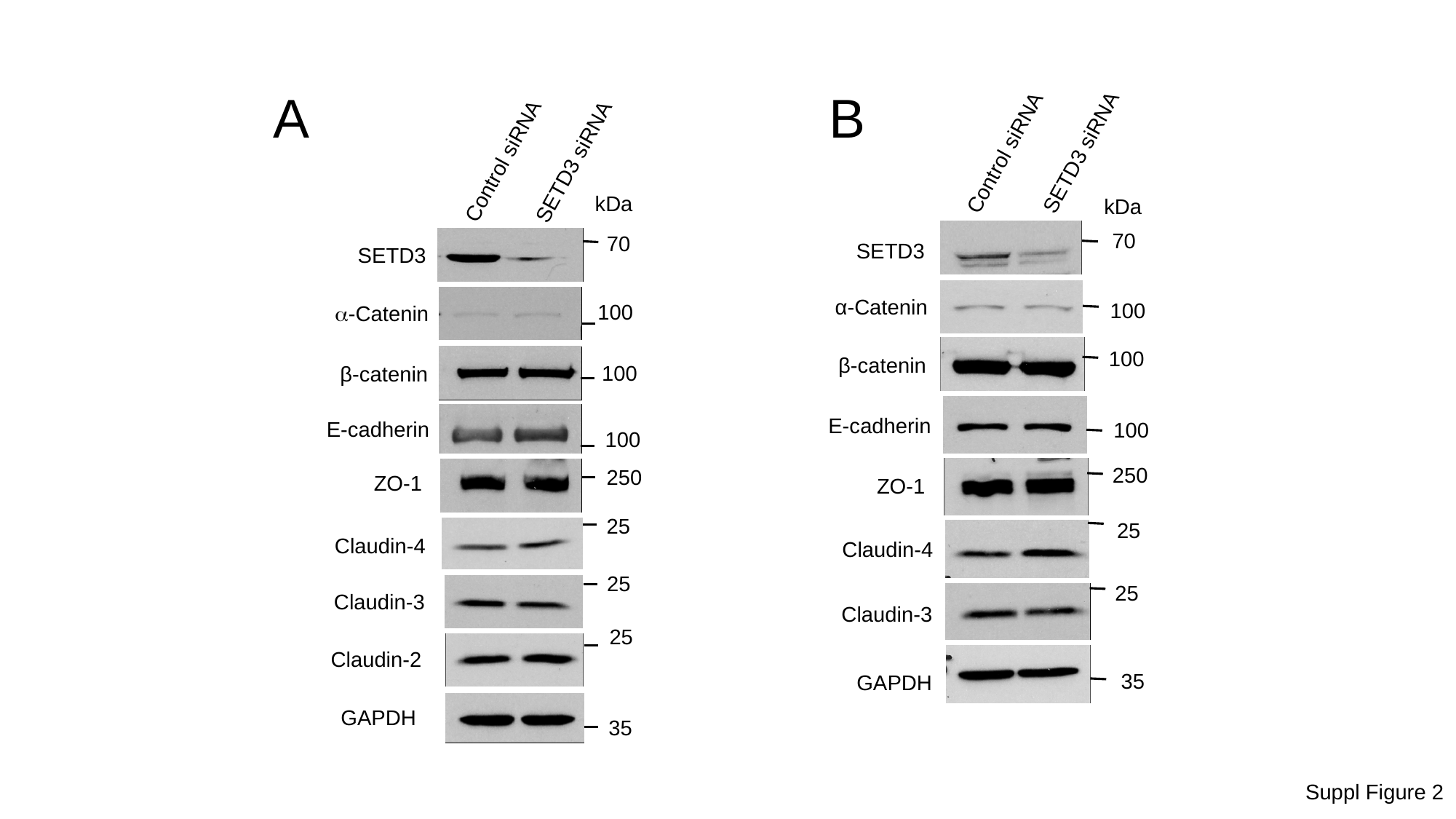

B
A
Control siRNA
SETD3 siRNA
Control siRNA
SETD3 siRNA
kDa
kDa
70
70
SETD3
SETD3
α-Catenin
100
100
α-Catenin
100
β-catenin
100
β-catenin
E-cadherin
E-cadherin
100
100
250
250
ZO-1
ZO-1
25
25
Claudin-4
Claudin-4
25
25
Claudin-3
Claudin-3
25
Claudin-2
35
GAPDH
GAPDH
35
Suppl Figure 2
