## Supplemental figure3 for "SETD3 ACTIN HISTIDINE METHYLTRANSFERASE REGULATES INTESTINAL EPITHELIAL BARRIER INTEGRITY AND MUCOSAL INFLAMMATION"

### Slide 1
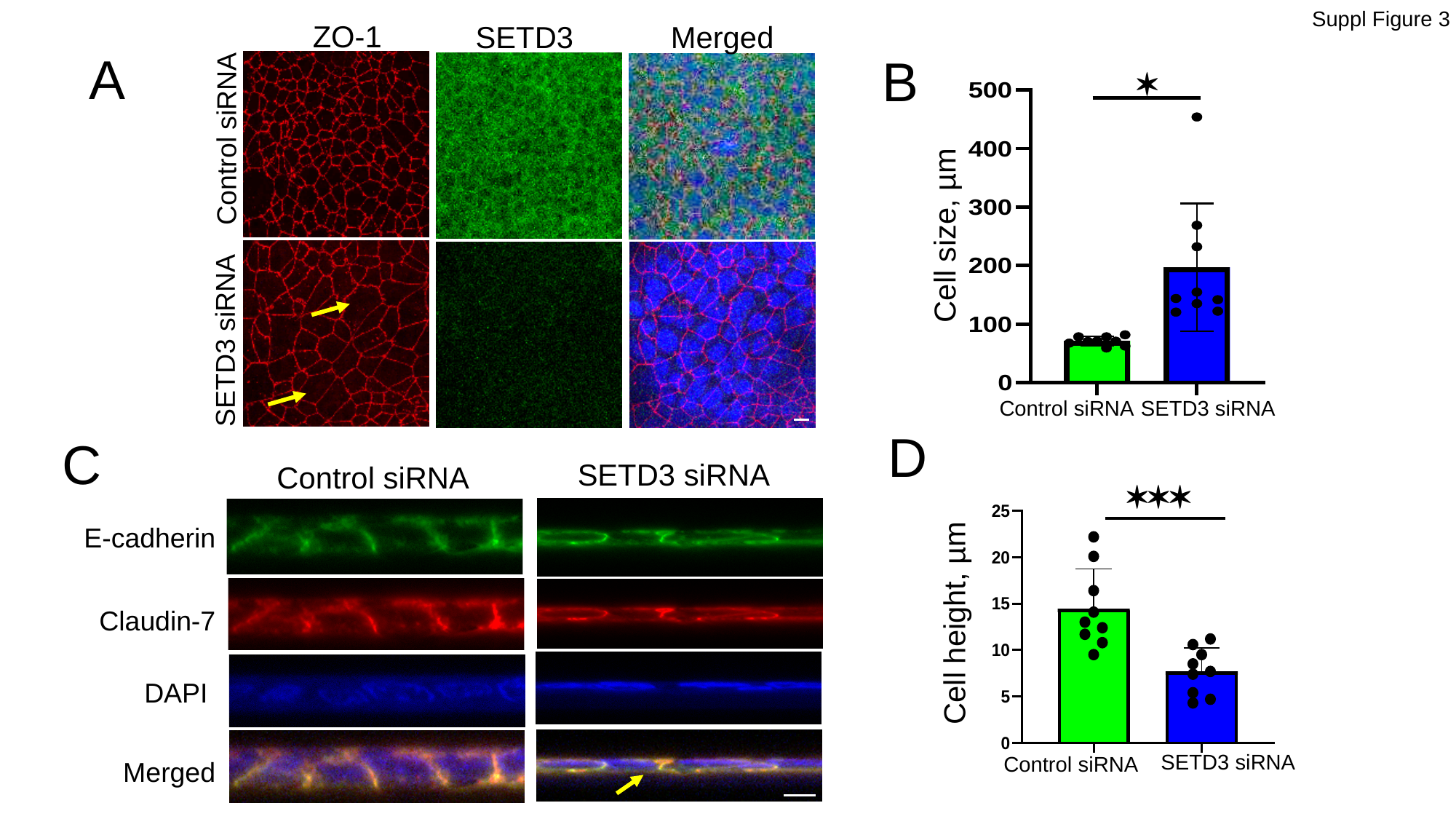

Suppl Figure 3
ZO-1
SETD3
Merged
A
B
*
Control siRNA
Cell size, µm
SETD3 siRNA
Control siRNA
SETD3 siRNA
D
C
SETD3 siRNA
Control siRNA
***
E-cadherin
Cell height, µm
Claudin-7
DAPI
SETD3 siRNA
Control siRNA
Merged
