## Supplemental figure4 for "SETD3 ACTIN HISTIDINE METHYLTRANSFERASE REGULATES INTESTINAL EPITHELIAL BARRIER INTEGRITY AND MUCOSAL INFLAMMATION"

### Slide 1
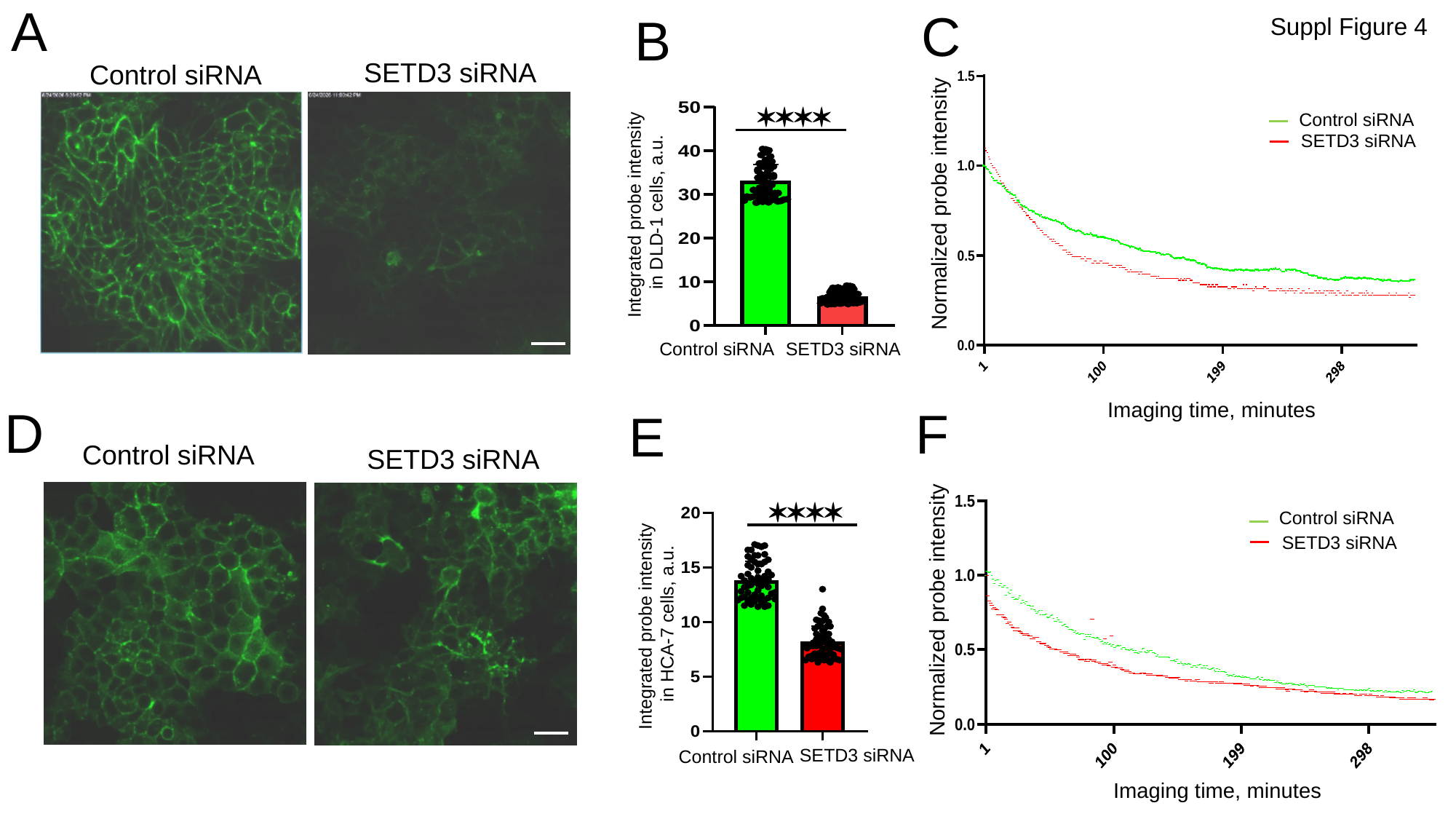

A
C
B
Suppl Figure 4
SETD3 siRNA
Control siRNA
****
Control siRNA
 SETD3 siRNA
Integrated probe intensity
in DLD-1 cells, a.u.
Normalized probe intensity
Control siRNA
SETD3 siRNA
Imaging time, minutes
D
F
E
Control siRNA
SETD3 siRNA
****
Control siRNA
 SETD3 siRNA
Normalized probe intensity
Integrated probe intensity
 in HCA-7 cells, a.u.
SETD3 siRNA
Control siRNA
Imaging time, minutes
