## Supplemental figure5 for "SETD3 ACTIN HISTIDINE METHYLTRANSFERASE REGULATES INTESTINAL EPITHELIAL BARRIER INTEGRITY AND MUCOSAL INFLAMMATION"

### Slide 1
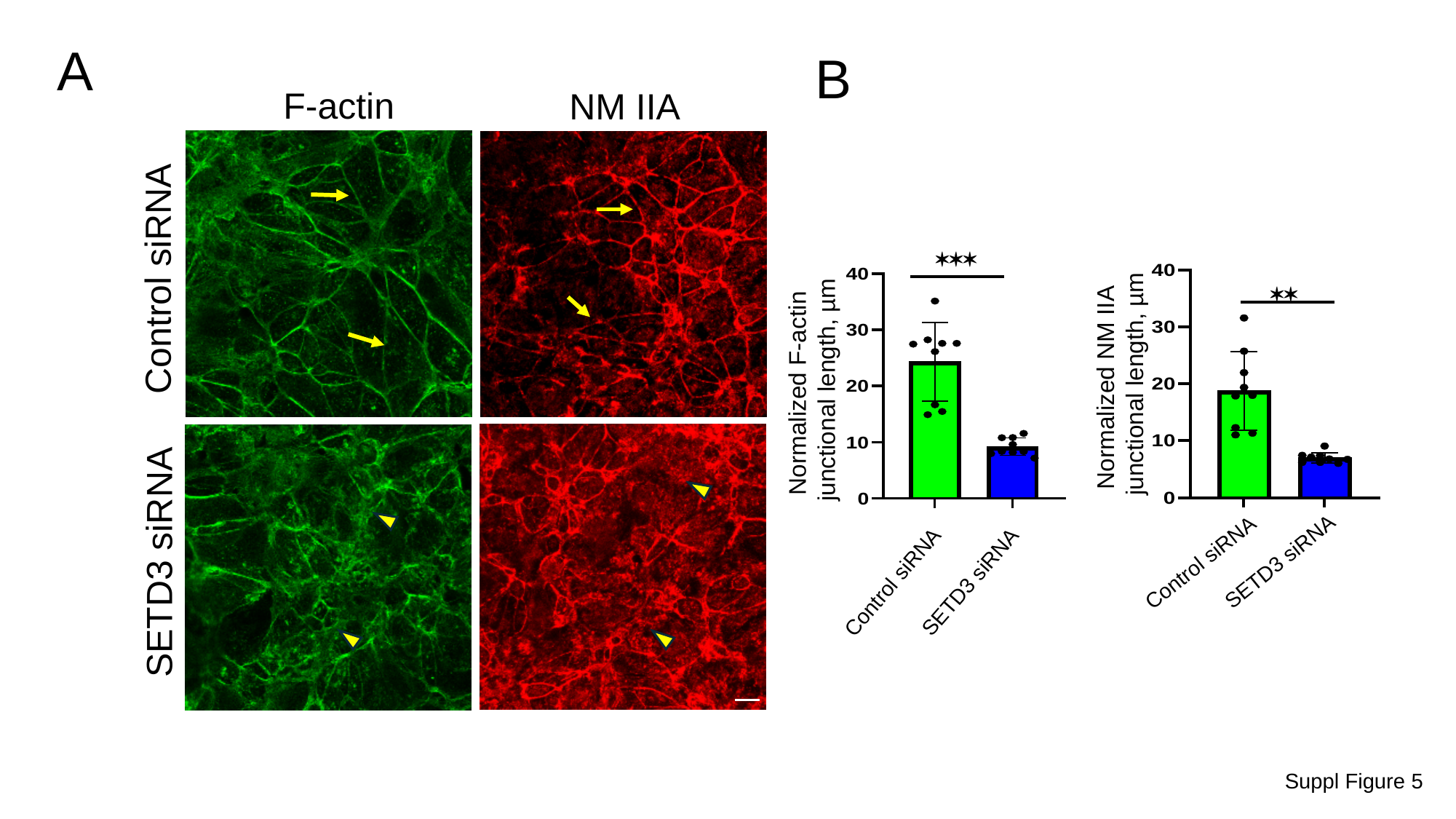

A
B
F-actin
NM IIA
***
Control siRNA
**
Normalized NM IIA
 junctional length, µm
Normalized F-actin
junctional length, µm
SETD3 siRNA
SETD3 siRNA
Control siRNA
Control siRNA
SETD3 siRNA
Suppl Figure 5
