## Supplementary figures and images for "SETD3 ACTIN HISTIDINE METHYLTRANSFERASE REGULATES INTESTINAL EPITHELIAL BARRIER INTEGRITY AND MUCOSAL INFLAMMATION"

### Supplemental figure6

#### Slide 1
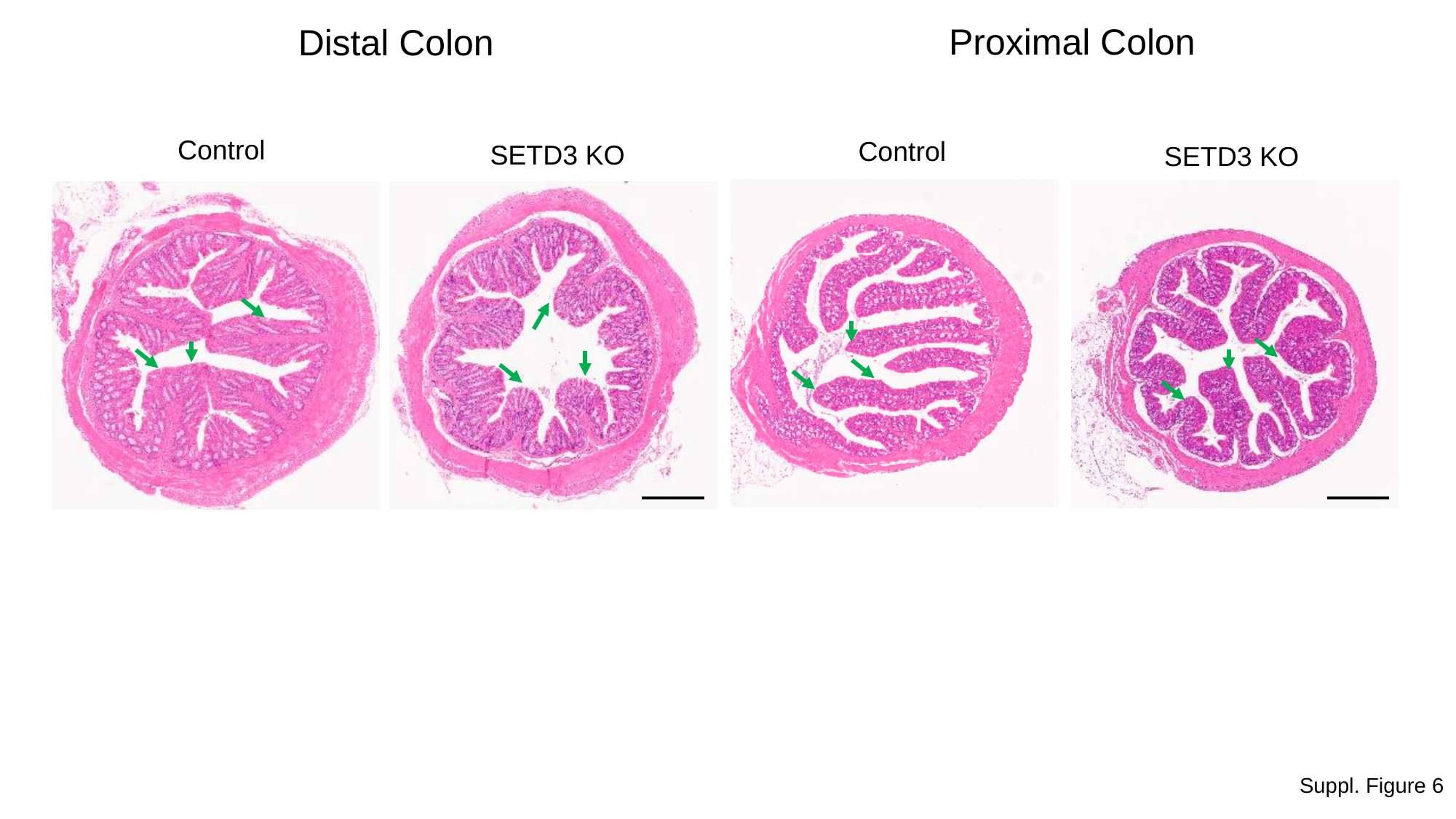

Proximal Colon
Distal Colon
Control
Control
SETD3 KO
SETD3 KO
Suppl. Figure 6
