## Supplemental figure7 for "SETD3 ACTIN HISTIDINE METHYLTRANSFERASE REGULATES INTESTINAL EPITHELIAL BARRIER INTEGRITY AND MUCOSAL INFLAMMATION"

### Slide 1
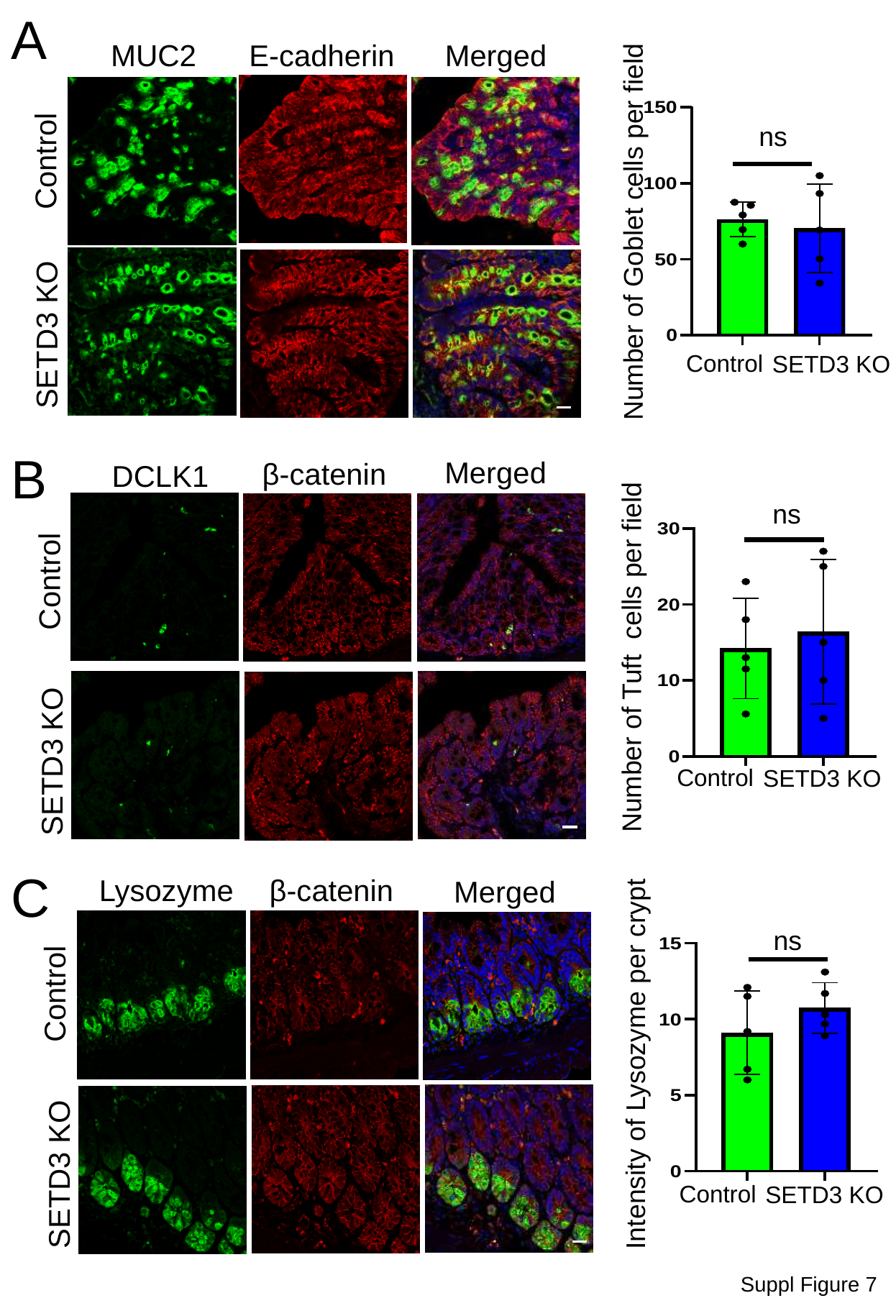

A
MUC2
Merged
E-cadherin
ns
Control
Number of Goblet cells per field
SETD3 KO
Control
SETD3 KO
B
Merged
β-catenin
DCLK1
ns
Control
Number of Tuft cells per field
SETD3 KO
Control
SETD3 KO
C
Lysozyme
β-catenin
Merged
ns
Control
Intensity of Lysozyme per crypt
SETD3 KO
Control
SETD3 KO
Suppl Figure 7
